# Single-Cell Transcriptome Analyses Reveal the Cell Diversity and Developmental Features of Human Gastric and Metaplastic Mucosa

**DOI:** 10.1101/2022.05.22.493006

**Authors:** Ayumu Tsubosaka, Daisuke Komura, Hiroto Katoh, Miwako Kakiuchi, Takumi Onoyama, Asami Yamamoto, Hiroyuki Abe, Yasuyuki Seto, Tetsuo Ushiku, Shumpei Ishikawa

## Abstract

The stomach is an important digestive organ with a variety of biological functions. However, due to the complexity of its cellular and glandular composition, the precise cellular biology has yet to be elucidated. In this study, we conducted single-cell RNA sequence analysis of the human stomach and constructed a 137,610-cell dataset, the largest cell atlas reported to date. By integrating this single-cell analysis with spatial cellular distribution analysis, we were able to clarify novel aspects of the developmental and tissue homeostatic ecosystems in the human stomach. We identified *LEFTY1*+ as a potential stem cell marker in both gastric and intestinal metaplastic glands. We also revealed skewed distribution patterns for PDGFRA+BMP4+WNT5A+ fibroblasts that play pivotal roles in, or even precede, the phenotypic changes from gastric to metaplastic mucosa. Our extensive dataset will function as a fundamental resource in investigations of the stomach, including studies on development, aging, and carcinogenesis.

## INTRODUCTION

The stomach is an essential digestive organ found in many organisms. Its roles include storing and digesting food, releasing food to the intestine, secreting various digestive enzymes, and releasing hormones, e.g., gastrin and somatostatin (Voutilainen et al., 2002). Additionally, mucosal immunity is prevalent in the stomach through exposure to organisms/molecules swallowed during daily life (Nie and Yuan, 2020). The stomach mucosa consists of epithelial glands with a variety of compositions and functions that are biologically and histologically classified into three subtypes. One of the signature glands of the stomach, the fundic gland, is found in the fundus/corpus and composed mainly of chief cells, parietal cells, endocrine cells, and mucous cells. Another stomach-specific gland, the pyloric gland (PG), is found in the pylorus and composed mainly of mucous cells and endocrine cells. The other gland found in the stomach, the intestinal metaplasia (IM) gland, mimics the colorectal epithelial crypt and is a metaplastic gland associated with atrophy and caused by chronic inflammation, such as that resulting from *Helicobacter pylori* infection (Wroblewski et al., 2010).

Each stomach gland has a dedicated stem cell source (Kim and Shivdasani, 2016); however, in contrast to other digestive organs, such as the esophagus and intestines, the high cell diversity and complexity of the stomach has led to difficulty in identifying stomach epithelial stem cells related to developmental biology. Thus, similarities and differences in the developmental properties of the three gastric gland subtypes remain to be investigated. Although several stomach epithelial or pan stem cell markers, such as LGR5, CD44 (Kim and Shivdasani, 2016; Ye et al., 2018), and AQP5 (Tan et al., 2020), have been proposed, a consensus on such markers has yet to be reached. The identification of stem cells is important if we are to understand the development of tissues and tumorigenesis. Notably, IM of the gastric mucosa is a well-known pathological condition that results directly in gastric carcinoma (Wroblewski et al., 2010); therefore, it is important to clarify how metaplastic mucosa arise from otherwise healthy gastric mucosa. Gastric stem cells may be transformed into intestinal stem cells (Jang et al., 2015; Simmini et al., 2014); however, the precise developmental properties of gastric and metaplastic glands have yet to be determined. Moreover, the differences and/or similarities between intestinal metaplastic gastric mucosa and genuine colorectal mucosa have not been clarified. Cell–cell communication through signaling molecules, such as cytokines, chemokines, and growth factors, is fundamental to establishing appropriate local tissue homeostasis in the human body. IM of the stomach can be triggered by pathological cycles of chronic inflammation and tissue repair; thus, the roadmap to IM may be affected by cellular communication in the environment of regenerative gastric mucosa, including that involving epithelial cells and various stromal cells, such as fibroblasts.

With the rapid development of single-cell RNA sequencing (scRNA-seq) and its associated analytical methods (Hao et al., 2021; Stuart et al., 2019), it has become feasible to construct global cell atlases with single-cell resolution and employ these to perform detailed analyses, e.g., identifying potential stem cell populations, clarifying developmental trajectories, and determining cell–cell communications between given cell types. To date, even the largest single cell atlas of the whole human body (The Tabula Sapiens Consortium, 2022) has not included the stomach scRNA-seq dataset. Busslinger et al. (2021) reported a scRNA-seq profile of human upper gastrointestinal tract; however, a specific stomach scRNA-seq dataset has been lacking.

In the present study, we constructed the largest ever transcriptional cell atlas of adult human gastric tissues using scRNA-seq and used it for the analyses as follows: profiling the global cellular diversity of the complexed gastric mucosa in healthy and metaplastic tissues, identifying possible stem cell populations in gastric glands, and discovering novel cell–cell communications related to homeostasis in the studied conditions. By integrating sophisticated bioinformatics analysis of scRNA-seq data with high-resolution spatial distribution analyses of specific mRNA molecules in human tissues, we not only identified a possible stem cell cluster common among the gastric glands but also clarified the spatially and functionally defined biological roles of BMP4-secreting fibroblasts found in either healthy or metaplastic gastric mucosa. As a resource that includes a scRNA-seq dataset with 137,610 cells, our human stomach cell atlas could help researchers provide new insights in the fields of gastric development, gastric stem cell biology, and gastric carcinogenesis.

## RESULTS and DISCUSSION

### Single-cell atlas of normal and IM gastric mucosa

We obtained gastric tissues from 15 patients who underwent gastrectomy at The University of Tokyo Hospital, and their scRNA-seq data were combined with those of 9 patients from Stanford University (Sathe et al., 2020) and 9 patients from Tsinghua University (Zhang et al., 2019). These nontumor gastric tissues were derived from a spectrum of healthy and disease states, including histologically normal gastric tissues, gastric cancers, intestinal metaplastic mucosa, and gastritis specimens (Table S1).

After exclusion of low quality and doublet cells, 137,610 cells were retained (see Methods; Figure 1A). After batch effect correction, unsupervised clustering analysis was used to identify 35 clusters. We merged the clusters into seven major cell lineages based on differential gene expression as follows (Figures 1B and 1C): 39,169 epithelial cells (characterized by *KRT19, TFF1,* and *PGA4*), 71,360 B and plasma cells (B cells: *MS4A1*; plasma cells: *IGHG1, IGHA1, IGKC,* and *IGHG4*), 15,778 T cells (*CD3D*), 2,002 myeloid cells (*FCGR3A* and *ITGAM*), 5,225 fibroblasts (*COL1A1* and *ACTA2*), 3,071 endothelial cells (*PECAM1* and *VWF*), and 1,005 mast cells (*TPSAB1*). The proportions of cell types in each clinical procedure or institution are shown in Figure 1D. Some of the samples in our institution contained exclusively higher proportions of B and plasma cells; thus, we excluded these samples from this research. Biopsy specimens from Tsinghua University had substantially larger proportions of epithelial cells, consistent with the fact that gastric biopsies mainly obtain surface mucosa. Additionally, surgical specimens from Stanford University and our institution had a larger proportion of nonepithelial cells. Overall, by combining three large RNA-seq datasets of the stomach with different clinical procedures and disease states, we successfully obtained a well-balanced and diverse cell atlas of the human stomach.

**Figure 1.**
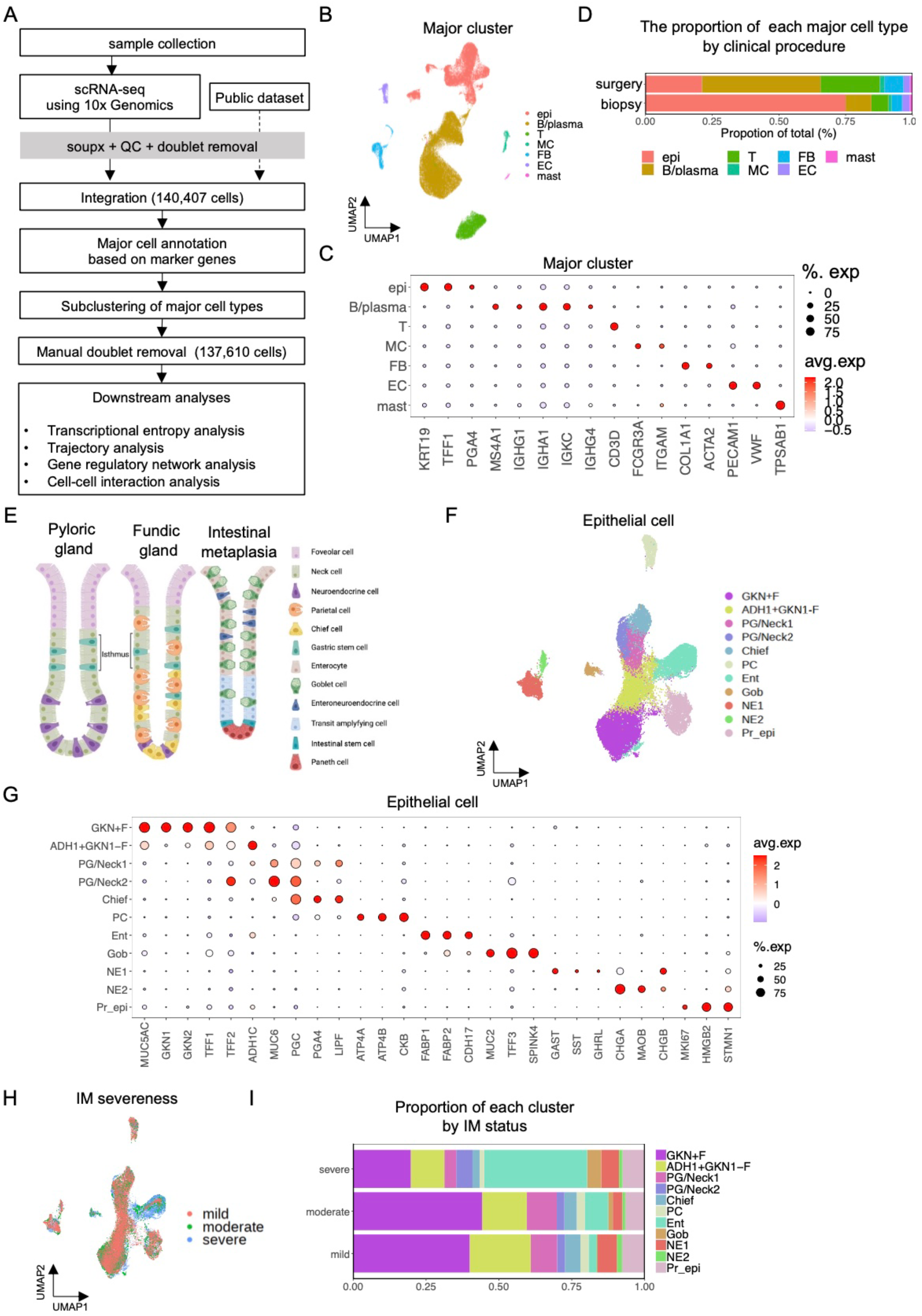
Major cell and epithelial cell clustering in stomach mucosa. (A) Overview of the comprehensive scRNA-seq analysis workflow. (B) UMAP showing the 7 major cell types identified by scRNA-seq (137,610 cells) of all cells after QC. (C) Representative major cell marker genes. Size and color of circles show the percentage of cells expressing genes and average gene expression, respectively. (D) Proportion of major cell types according to clinical procedure. Biopsy specimens had larger proportions of epithelial cells and surgical specimens had a larger proportion of nonepithelial cells. (E) Scheme of each epithelial cell-type in the pyloric gland, fundic gland, and metaplastic mucosa. (F) UMAP showing subclusters of epithelial cells (39,169 cells) identified by scRNA-seq. (G) Representative epithelial cell marker genes. Size and color of circles show the percentage of cells expressing genes and average gene expression, respectively. (H) UMAP showing the IM state tissue from which each cell was derived. See methods for IM status determination. (I) Proportion of epithelial cell types according to IM status. IM severe samples had a larger proportion of goblet and enterocytes. Abbreviations: UMAP, Uniform Manifold Approximation and Projection; QC: quality check; epi: epithelial cells; MC: myeloid cells; FB: fibroblasts; EC: endothelial cells, F: foveolar cells, PG: pyloric gland cells, PC: parietal cells, Ent: enterocytes, Gob: goblet cells, NE: neuroendocrine cells, Pr_epi: proliferating epithelial cells.

### Epithelial cells

#### Subcluster determination

We subclustered 39,169 epithelial cells into 11 clusters comprising foveolar subtypes (characterized by *MUC5AC, GKN1, GKN2, TFF1, TFF2,* and *ADH1C*), PG/neck cells (*MUC6, PGC,* and *TFF2*), chief cells (*PGA4, PGC,* and *LIPF*), parietal cells (*ATP4A, ATP4B,* and *CKB*), enterocytes (*FABP1, FABP2,* and *CDH17*), goblet cells (*MUC2, TFF3,* and *SPINK4*), neuroendocrine (NE) cells (*GAST, SST, GHRL, CHGA, MAOB,* and *CHGB*), and proliferating cells (*MKI67, HMGB2,* and *STMN1*) (Figures 1E–G; Figure S1A). The proportion of cells in each subcluster along with the severeness of IM is shown in Figures 1H and 1I, showing the increase in the number of enterocytes and goblet cells in the severe IM samples.

First, we focused on the characteristics of NE cells because they might represent the interpretable characteristics of each gastric and metaplastic gland. NE cells commonly expressed *CHGA* and *CHGB* and were clustered into at least five populations, including G cells and D cells, based on the expression of hormones or enzymes, such as *SST* (D cell marker)*, GHRL* (X/A-like cell marker), *GAST* (G cell marker), and *MAOB* (enterochromaffin cell marker; Figures S1B and S1C; Busslinger et al. (2021)). Some NE cells expressed *LHB*, as reported by Busslinger et al. (2021). Interestingly, some NE cells expressed *REG4*, a specific marker of metaplastic mucosa (Zhang et al., 2019). Some *REG4*+ NE cells also expressed *GCG* and *PYY*, which are specific to enteroneuroendocrine cells (Gunawardene et al., 2011). Notably, the severeness of IM (Figure 1I) was correlated with the frequency of *REG4*+ NE cells (Figures S1B and S1C), suggesting that IM of the stomach consists of NE cells with the endocrine features of genuine colonic enteroneuroendocrine cells. The phenotypic similarity between IM of the stomach and colonic mucosa indicates that the regeneration of atrophic gastric mucosa generates metaplastic mucosa that resembles, both histologically and functionally, genuine colorectal mucosa.

Foveolar cells, surface mucous cells for which the biological characteristics are known to differ between gastric and metaplastic mucosa (Kim and Shivdasani, 2016), were clustered into two distinct populations: GKN1+F cells and ADH1+GKN1-F cells (Figures 1F and 1G; Figure S1A). Encoded by *GKN1*, gastrokine-1 is a stomach-specific protein with various functions, including modulating cell cycle progression, cellular proliferation, and antibiotic, anti-inflammation, and antiapoptotic actions (Alarcón-Millán et al., 2019). Encoded by *ADH1C*, alcohol dehydrogenase 1C is often discussed in the context of ethanol metabolism (Edenberg and McClintick, 2018); however, the relationships between its expression and *H. pylori* infection and IM have been investigated previously, and it may be relevant to the metabolism of retinol acid (Matsumoto et al., 2005). In human stomach specimens, GKN1 expression was observed in gastric mucosa but not in metaplastic mucosa (Figures S1D and S1E); moreover, the spatial distributions of the distinct GKN1+F and ADH1+GKN1-F populations showed clear gradation patterns in the superficial and deeper layers, respectively, of the gastric mucosa (Figures S1D and S1E). Thus, a combination of scRNA-seq analysis and spatial identification of specific populations confirmed the histological distributions of the distinct foveolar subtypes, consistent with the findings of previous studies (Mao et al., 2012; Westerlund et al., 2007); moreover, the results suggested that the ADH1+GKN1-F and GKN+F populations represent immature (deeper layer) and mature (surface layer) foveolar epithelium, respectively.

The transcription factor *NKX6-3* is known to be a distinctive positive modulator of *GKN1* (Alarcón-Millán et al., 2019), and its expression is specific to the gastric mucosa (Choi et al., 2008). NE cells showed characteristically higher expression of *NKX6-3*, especially G and D cells (Figures S1A and S1C). We also found that non-NE cells in the stomach showed various degrees of positivity for *NKX6-3* (Figure S1A), indicating that *NKX6-3* has a wider range of biological functions in various stomach cells than was previously expected. Indeed, a previous study showed that *NKX6-3* inactivation in the stomach led to overexpression of *CDX2* and reduced expression of *SOX2* (Yoon et al., 2015).

Gastric fundic glands and PGs are composed of foveolar epithelium, isthmus, and neck cells (Figures 1E and 1F). Using our integrated scRNA-seq dataset, we identified diverse cell populations, including those of the neck areas of gastric glands, which we termed PG/Neck cells (Figures 1E–G). Intriguingly, we identified two distinct PG/neck cell populations: PG/Neck1 and PG/Neck2 cells (Figures 1F and 1G). Some PG/Neck2 cells expressed *MUC6* and/or *TFF2*; thus, they included pyloric as well as fundic mucous gland cells (Wuputra et al., 2021; Zhang et al., 2019). Higher expression of *MUC6* and *TFF2* is reportedly related to spasmolytic polypeptide-expressing metaplasia (Nam et al., 2010), which is associated with chronic inflammation and IM of the stomach (Radyk et al., 2018). Consistent with this notion, the PG/Neck2 population included cells that expressed *CLDN2* and *TFF3*, known markers for intestinal and goblet cells, respectively (Escaffit et al., 2005; Zhang et al., 2019). Thus, the PG/Neck2 cells apparently include highly diverse cell populations covering the fundic/pyloric and metaplastic glands. These cells also expressed *AQP5*, *ODAM*, and *PRR4* as differentially expressed markers, all of which are known salivary gland markers (Hosoi, 2016; Huang et al., 2021), although the underlying physiological basis of the similarity between the stomach and saliva is unclear. Notably, *AQP5* was proposed as a gastric stem cell marker in a previous study (Tan et al., 2020).

### Transcriptional entropy and gene expression trajectory analyses of epithelial cells reveal the LEFTY1+ cell population as a potential stem cell cluster common to the gastric and metaplastic glands

The distinct stem cell populations in the gastric and metaplastic mucosa are yet to be clarified; however, it is hypothesized that the stem cell compartment exists in the PG/Neck cells among the cellular clusters identified here (Han et al., 2019; Kim and Shivdasani, 2016). To determine the possible stem cells of the stomach glands, we performed transcriptional entropy analysis, calculating the stemness score based on the number of expressed genes per cell (see Methods; Gulati et al., 2020). The stemness score was considered an indicator of the differentiation states of each cell: higher entropy suggests that the cell is in a more immature state (Gulati et al., 2020). In our analysis, PG/Neck2 cells had significantly higher stemness scores compared with those of PG/Neck1 cells (p<2.2e-16, Figure 2A); therefore, we hypothesized that the PG/Neck2 cluster contained possible stem cells. To explore the stem cell populations of the stomach epithelial cells as well as their developmental paths, we performed unsupervised trajectory analysis (Cao et al., 2019) in which the differentiation dynamics of gene expression were visualized (Figures 2B–E). This analysis revealed two separated lineages, namely the normal gastric lineage and the intestinal metaplastic lineage, which were characterized by *SOX2* and *CDX2* enrichment, respectively (Figures 2B and 2C), consistent with their known functions in the respective development of the stomach and intestine (Kim and Shivdasani, 2016). Interestingly, we found a tiny population in PG/Neck2 cells (we termed these “linking cells”) located between the gastric and intestinal routes (Figure 2D). In the PG/Neck2 population, these distinct cells were thought to be possible stem cells because expression of the stem-associated markers (e.g., *AQP5, LGR5, SMOC2, ASCL2, TNFRSF19, EPHB2, CD44,* and *PROM1*; Guo and Frenette, 2014; Jang et al., 2013, 2017; Kim and Shivdasani, 2016; Tan et al., 2020; Ye et al., 2018) was relatively high (Figures S2A and S2B). Moreover, pseudotime analysis showed chronological trajectories from the “linking cell” population to various paths of the stomach epithelia (Figure 2E). Among PG/Neck2 cells, the candidates of specific markers for “linking” cells were *LEFTY1*, *OLFM4*, and *CLDN4* in differentially expressed genes (Figure 2F). Whereas *OLFM4* and *CLDN4* were expressed not only in linking cells, but in the enterocytes and goblet cells (Figures S2A and S2B), LEFTY1 was highly enriched in the “linking” cells (Figure 2G).LGR5, a representative stem cell marker, was also expressed in the “linking” cells (Figure 2G). Above all, *LEFTY1* was distinguished as a differentially expressed gene in the “linking” cells with possible stem cell properties.

**Figure 2.**
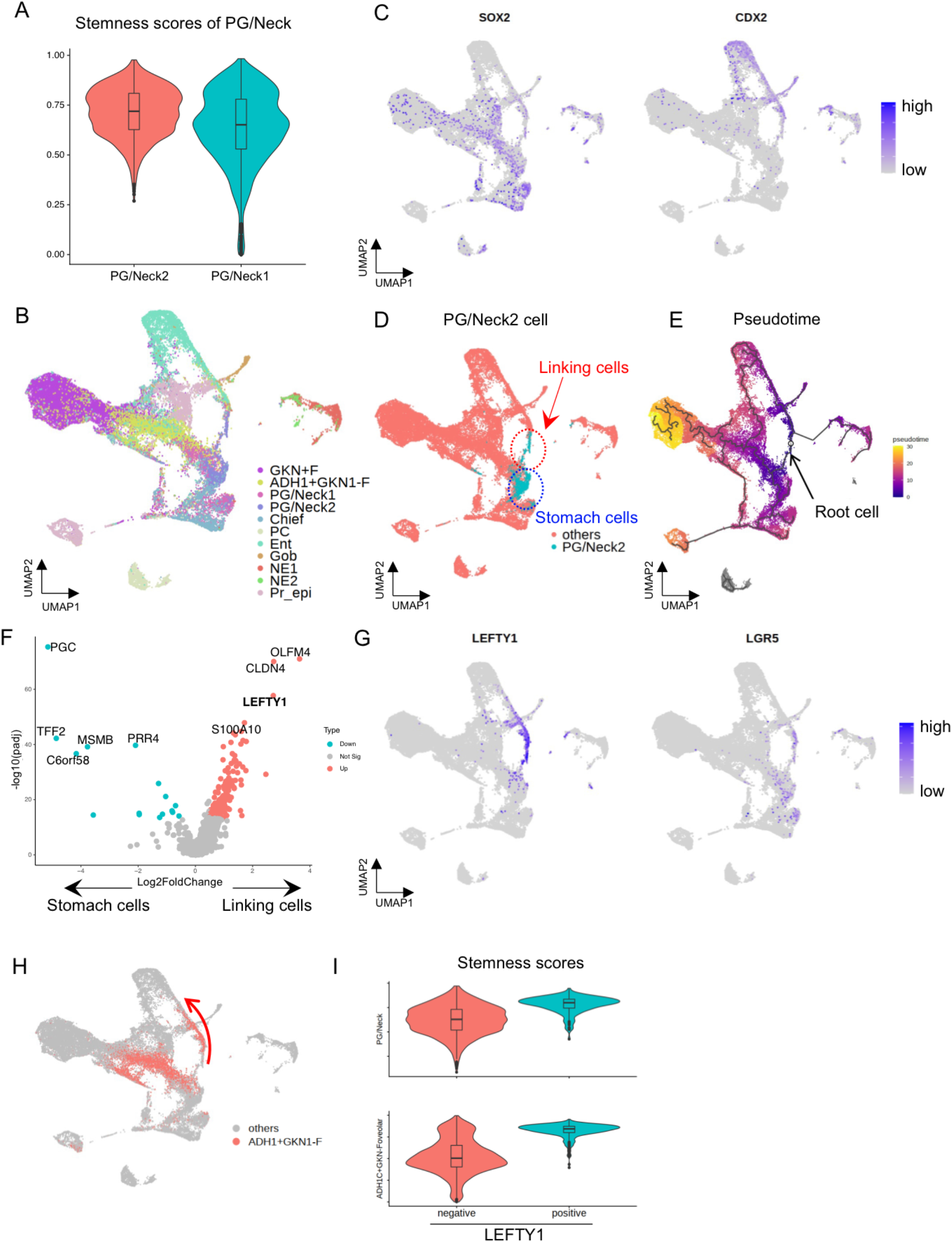
Trajectory analysis of epithelial cells identified a possible novel stem cell marker, LEFTY1. (A) The comparison of stemness scores between PG/Neck1 and 2. The score of PG/Neck2 is significantly higher than PG/Neck1 (p<2.2e-16, two-sided Welch’s t test). (B) UMAP showing each subcluster shown in Figure 1F. UMAP coordinates of epithelial cells was re-calculated by trajectory inference analysis. See Methods for details. (C) *SOX2* and *CDX2* expression (master regulators of the stomach and intestine, respectively) in epithelial cells. *SOX2* was expressed in the normal gastric cells, whereas *CDX2* was in the intestinal metaplastic cells. (D) UMAP showing two separate groups of PG/Neck2 cells: linking cells, which were located between the metaplastic and stomach cells, and those within the other stomach cells. (E) Pseudotime trajectory analysis displayed on the UMAP plot. It is based on the assumed root cell (arrow), which was determined manually among LEFTY1+ positive cells. (F) Volcano plot showing the top differentially expressed genes between the two PG/Neck2 groups shown in Figure 2D (linking cells and stomach cells). *OLFM4*, *CLDN4*, and *LEFTY1*, were top differentially expressed genes in the linking cells. (G) *LEFTY1* and *LGR5* expression in epithelial cells. Both genes were expressed in the linking cells. (H) ADH1+GKN1-F cells on the UMAP plot. Some ADH1+GKN1-F cells on the routes of metaplastic lineages (arrow) expressed *LEFTY1* as shown in Figure 2G. (I) The comparison of stemness scores between LEFTY1+ and LEFTY1− cells in each PG/Neck2 cell cluster and ADH1+GKN-F cell cluster. The score of each LEFTY1+ population is significantly higher than LEFTY1-population (p<2.2e-16, two-sided Welch’s t test). Abbreviations: padj, adjusted p-value.

*LEFTY1*, the product of which is a secreted protein and transforming growth factor-beta (TGF-β) superfamily member, has been extensively studied in the developmental stage and is known to play a role in determining left–right asymmetry (Kosaki et al., 1999; Meno et al., 1998). LEFTY1 inhibits SMAD signaling by binding to Cripto-1 and blocks Nodal in the development of mice (Tabibzadeh and Hemmati-Brivanlou, 2006). Additionally, scRNA-seq analysis of Barrett’s esophagus has shown that *LEFTY1* is a potential marker of Barrett’s esophagus precursors in human (Owen et al., 2018). Zabala et al. (2020) showed that LEFTY1 and bone morphogenetic protein (BMP) 7 maintained long-term proliferation and differentiation of human mammary gland cells, respectively, through a mechanism whereby LEFTY1 binds to BMPR2 and prevents BMP7/BMPR2-mediated SMAD activation. In the present study, some ADH1+GKN1-F cell routes specifically found in the metaplastic lineage also expressed *LEFTY1* (Figures 2G and 2H). To determine whether *LEFTY1* is an actual marker of stem cells and investigate their possible roles in the development of stomach mucosa, we divided both PG/Neck2 and ADH1+GKN1- F cells into two clusters each, i.e., *LEFTY1*+ and *LEFTY1−*, respectively. First, *LEFTY1*+ cells showed significantly higher stemness scores in both PG/Neck2 and ADH1+GKN1- F cell populations (p*<*2.2e-16, Figure 2I). Pseudotime plotting showed that *LEFTY1* was highly expressed in cells at the earliest time point and commonly in all types of stomach glands (Figures S2C–H). Notably, some conventional stemness-associated genes, such as *CD44* and *EPHB2*, were also expressed in similar time courses to those in which *LEFTY1* was expressed (Figures S2C–E). In the middle of pseudotime, cell division marker, *MKI67*, were expressed around the “proliferative epithelial” population (Figures S2C–E). Compared with other conventional stem cell markers or with stem cell- or cancer stem cell-enriched markers (e.g., *AQP5, LGR5, SMOC2, ASCL2, TNFRSF19, EPHB2, CD44,* and *PROM1*; Guo and Frenette, 2014; Jang et al., 2013, 2017; Kim and Shivdasani, 2016; Tan et al., 2020; Ye et al., 2018), LEFTY1*+* cells were more highly expressed and specifically existed in a “linking” cell population (Figure 2G; Figures S2A and S2B). Cell cycle analysis showed that the ratio of G2/M cells was lowest in *LEFTY1*+ PG/Neck cells (Figure S2I), which was compatible with their quiescence.

IHC showed that LEFTY1+ cells existed in gastric pyloric and fundic glands at very low frequencies and were spatially located mainly around the so-called isthmus regions (Figure 3A), consistent with the consideration of the histological isthmus region as a stem cell zone (Figure 1E; Han et al., 2019; Kim and Shivdasani, 2016). Contrastingly, in the intestinal metaplastic mucosa, LEFTY1+ cells were observed at much higher frequencies and were spatially located at the base of crypts (Figure 3A), consistent with the knowledge that intestinal stem cells reside at crypt bases (Spit et al., 2018). These spatial data support the hypothesis that *LEFTY1* is a novel marker of gastric stem cells. LEFTY1+ cell frequencies were highest in the metaplastic glands, followed by the PGs and fundic glands, respectively (Figure 3A). LEFTY1 staining showed two different patterns: a moderate cytoplasmic staining pattern and an intense dot signal pattern. However, the functional differences, if any, of LEFTY1 in relation to these staining patterns are not clear, as reported previously in an esophageal study (Owen et al., 2018).

**Figure 3.**
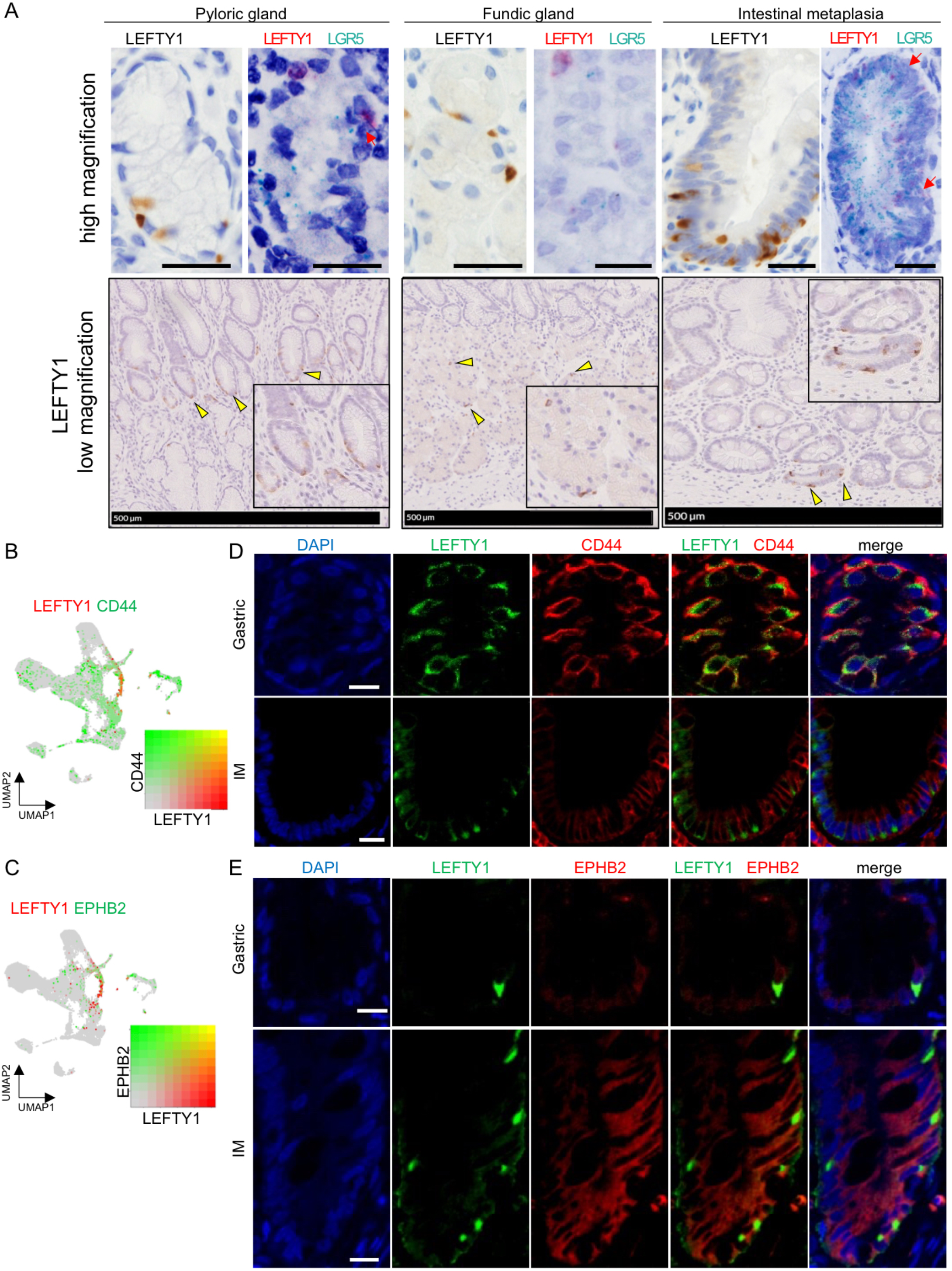
Coexpression of LEFTY1 and conventional gastrointestinal stem cell marker genes in epithelial cells. (A) Top: IHC of LEFTY1 (lefthand boxes) and RNA-ISH [righthand boxes: LEFTY1 (red) and LGR5 (green)]. Staining and *in situ* hybridization were performed in pyloric glands, fundic glands, and intestinal metaplasia (as shown). Some cells coexpressed LEFTY1 and LGR5 (arrows). Scale bar: 25 μm. Bottom: LEFTY1 low magnification of IHC. LEFTY1 expression was observed in the conventional stem cell zone. Arrowheads: LEFTY1+ cells. (B, C) Combined feature plots showing CD44 and EPHB2 (green) expression with LEFTY1 (red). Coexpressing cells are shown in yellow. (D, E) Immunofluorescence of LEFTY1 (green) and CD44/EPHB2 (red) in the intestinal metaplasia and gastric mucosa. (D) Area of clustered LEFTY1+ cells in gastric mucosa was selected for clarity. Scale bar: 10 μm.

We performed RNA *in situ* hybridization (RNA-ISH) of *LEFTY1* and *LGR5* (Figure 3A), finding that a portion of *LGR5*+ cells coexpressed *LEFTY1* in the pyloric and metaplastic mucosa. Additionally, scRNA-seq analysis and immunofluorescent staining showed that subsets of the CD44+ and/or EPHB2+ possible stem cells coexpressed LEFTY1 (Figures 3B–E). In our spatial analysis of human gastric tissues, the colocalization of LEFTY1 with other stem cell markers and the low frequency of LEFTY1+ cells among other stem-marker-positive cells strongly suggest that LEFTY1 is an actual candidate stem cell marker. LEFTY1+ cells can be considered common stem cells in both gastric and metaplastic glands based on our trajectory and pseudotime analyses; however, we found that EPHB2 was expressed in LEFTY1+ cells in the metaplastic mucosa but not in the normal gastric mucosa (Figure 3E). We hypothesize that, during IM, a phenotypic change occurs in LEFTY1+ stem cells in the normal gastric gland and they acquire the distinctive properties of intestinal stem cells by obtaining the EPHB2+ phenotype.

### Gene regulatory network analysis of stomach epithelial cells

To investigate the global gene regulatory network in gastric epithelial cells, we analyzed the activity of transcriptional programs in each epithelial cell-type by integrating the expression of transcription factors and their downstream target genes (Aibar et al., 2017; Van de Sande et al., 2020). Through gene regulatory network analysis, we obtained regulon activity scores in each cell-type (Figure 4A; Figure S3A). Our results were consistent with those of previous studies; for example, NE cells showed high paired box 6 (PAX6) or achaete-scute family BHLH transcription factor 1 (ASCL1) activities, as reported previously (Kim and Shivdasani, 2016), parietal cells showed high estrogen-related receptor gamma and beta activity, which is known to regulate ATP4A and ATP4B (Zhang et al., 2019), and enterocytes showed high scores for the colon-specific transcription factors CDX1 and CDX2 (Almeida et al., 2003) (Figure S3A). These results demonstrate the utility of gene regulatory network analysis for correctly identifying transcription factor activity.

**Figure 4.**
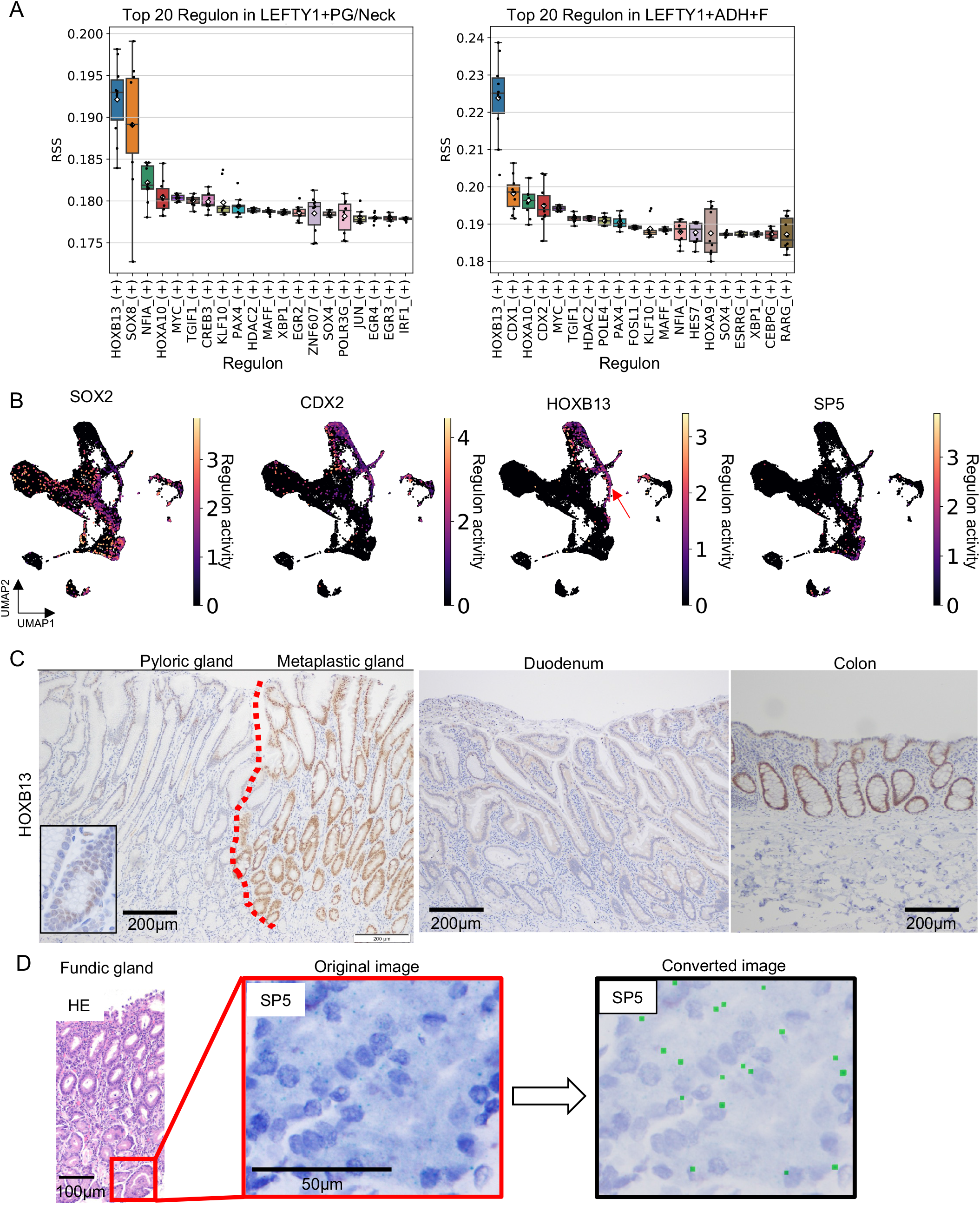
Gene regulatory network analysis in epithelial cells revealed LEFTY1+ cell-specific regulons. (A) LEFTY1+ cluster top 20 regulon specific scores calculated 10 times using pySCENIC. HOXB13 transcription factor showed highest scores commonly. LEFTY1+ADH+F showed relatively high scores of CDX1 and CDX2, suggesting these cells are on the routes of differentiation of cells destined for the metaplastic lineage. X-axis shows transcription factors. (B) UMAP plot showing transcription activities of SOX2, CDX2, HOXB13, and SP5. High HOXB13 regulon activity was observed in the LEFTY1+ cell population (arrow) and metaplastic cells. SOX2, CDX2, and SP5 regulon activity was high in the normal gastric cells, in the fundic gland specific cells, in the metaplastic lineage cells, respectively. (C) HOXB13 immunohistochemistry in the intestinal metaplasia, stomach, duodenum, and colon. HOXB13 expression was observed in the metaplastic gland and genuine colonic mucosa. No expression was observed in the duodenal mucosa. Inset: higher magnification of PG. Scale bar: 200 μm. (D) RNA-ISH showing SP5 expression in the base of the fundic gland. Right: larger green pixels converted computationally from original green signals (See methods for details).

Notably, high HOXB13 regulon activity was observed in LEFTY1+ clusters (in both LEFTY1+ PG/Neck and LEFTY1+ ADH+F clusters; Figure 4A). As expected, SOX2 and CDX2 regulon activities were high in normal and metaplastic mucosa, respectively (Figure 4B), whereas HOXB13 regulon activity was limited to the possible stem cell region and metaplastic epithelial cells (Figure 4B). Thus, the HOXB13 regulon might play an important role in the LEFTY1+ cells found in the metaplastic mucosa. *HOXB13*, the expression of which is almost exclusively found in the prostate and intestine, has been extensively studied in prostate cancer because both somatic and germline variants of *HOXB13* are associated with this cancer (Morgan and Pandha, 2017; Yu et al., 2020). HOXB13 is also known to downregulate the expression of TCF4 and its target MYC in colon cancer cells (Xie et al., 2019). Additionally, *HOXB13* expression is higher in left-sided colon cancers than that in right-sided colon cancers, and this higher expression level is associated with poor prognosis in right-sided colon cancers (Xie et al., 2019). In gastric cancer, HOXB13 promotes cell migration and invasion by upregulating PI3K/AKT/mTOR (Guo et al., 2021).

In the present study, IHC revealed high and universal HOXB13 expression in metaplastic mucosa, including in both complete and incomplete subtypes; however, in normal mucosa, protein expression was negligible (Figure 4C). These findings suggest that the HOXB13 regulon is indispensable in the development of IM and that the phenotypic switch of LEFTY1+ possible stem cells between gastric and metaplastic glands may require additional HOXB13 activation. Unlike in colonic mucosa, the expression of HOXB13 was not observed in duodenal or iliac mucosa, suggesting that metaplastic mucosa has similar characteristics to those of colonic mucosa (Figure 4C).

Regardless of *LEFTY1* expression, PG/Neck2 cells commonly showed high scores for SOX8 (Figure 4A; Figure S3A), for which an association with stomach biology has not been reported to date; thus, further investigation of its function in the stomach is warranted. Meanwhile, LEFTY1+ADH+F cells showed high scores for CDX1 and CDX2 (Figure 4A; Figure S3A), confirming that LEFTY1+ADH+F cells are on the routes of differentiation of cells destined for the metaplastic lineage (Figure 2H). Fundic gland-specific cells (i.e., chief and parietal cells) showed high SP5 activity (Figure S3A). Huggins et al. (2017) reported that SP5 is a WNT target and negatively regulates WNT activity in human pluripotent stem cells. We confirmed the specific expression of SP5 in fundic glands using RNA-ISH; thus, it appears to be important in the development and/or maintenance of these glands (Figure 4D; Figure S3B; see Methods).

Fazilaty et al. (2021) used scRNA-seq to show that embryonic enterocyte progenitor genes were reactivated in the damaged enterocytes of both humans and mice. Given that the stomach and intestine share common features, the gastric mucosa might also reactivate their progenitor programs upon epithelial damage. Indeed, we found that metaplastic mucosa expressed *LEFTY1* and *HOXB13*, both of which are embryonic genes (Kosaki et al., 1999; Ma et al., 2003), at higher levels than those in normal pyloric or fundic mucosa. Moreover, in our pseudotime analysis, mature metaplastic mucosa specifically expressed *APOA1*, one of the progenitor genes expressed in damaged colonic mucosa (Figure 2B; Figure S3C), indicating that metaplastic enterocytes were similar to immature colonic epithelium but not normal mucosa.

### Fibroblasts

As concluded by Higuchi et al. (2015), there are distinct differences in the stromal cells of the stomach and intestinal mucosa; therefore, specific fibroblasts play key roles in maintaining epithelial homeostasis in specific foci, and gastric fibroblasts are hypothesized to affect the developmental destinations of gastric or metaplastic epithelia in the stomach. Previous studies on colorectal and gastric glands have found that the proper compositions of fibroblasts and their secreting cytokines, including WNT, BMP, and TGF-β ligands/inhibitors, were fundamental to the development and maintenance of digestive tissue integrity (David et al., 2020; Koch, 2017; Wölffling et al., 2021; Zhang et al., 2020). In their scRNA-seq study of the whole human body, Buechler et al. (2021) showed that various organ-specific fibroblasts exist. However, global profiling of gastric-specific fibroblasts is lacking; thus, the characteristics of gastric fibroblasts, as well as their similarities and differences to those in metaplastic mucosa, remain to be elucidated. In the present study, we identified 5,225 fibroblasts, which were divided into 6 subclusters: KLF+ cells (characteristic expression of *KLF4, SFRP1, SFRP2,* and *PI16*), CCL11+ cells (*CCL11, ABCA8, APOE,* and *ADAM28*), PDGFR+ cells (*PLAT*, *POSTN*, *PDGFRA, BMP4,* and *WNT5A*), FibSmo (Fibroblasts which express both fibroblastic and smooth muscle markers; He et al., 2020) (*HHIP*, *MYLK*, and *ACTG2)*, smooth muscle cells (RERGL, ADIRF, MYH11, and *TAGLN*), and myofibroblasts (*RGS5* and *CD36*) (Figures 5A and 5B; Figure S4A).

**Figure 5.**
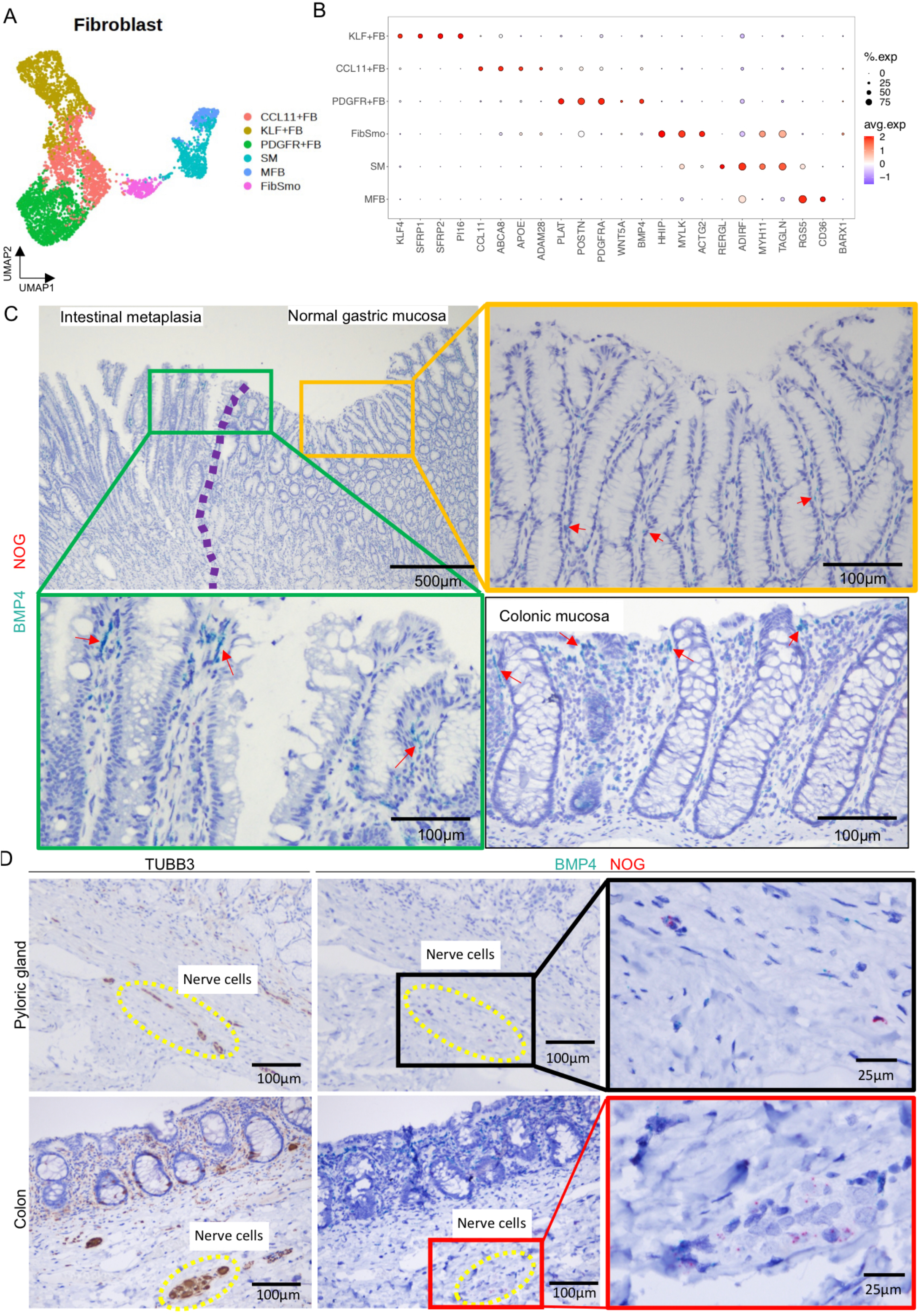
The increase of BMP4 expression in the fibroblasts that precedes the epithelial metaplastic transformation. (A) UMAP showing the subclusters of 5,225 fibroblasts. (B) Representative marker genes of fibroblast subclusters. Size and colors of circles show the percentage of cells expressing genes and average gene expression, respectively. (C) RNA-ISH of BMP4. Green contour shows IM and the transitional region; yellow contour shows the pyloric gland. BMP4 expression in IM and the transitional region was similar to that in genuine colonic mucosa. The increase of BMP4 expression was observed in the metaplastic gland and normal gastric mucosa adjacent to IM. No NOG expression was observed in mucosal laminar propria both in the gastric and colonic mucosa. Arrows: BMP4 signals. (D) Neuronal-specific TUBB3 IHC showing neuron cells in the submucosal region of the pyloric gland and colonic mucosa (Left). NOG expression was observed in the nerve cells both in the gastric and colonic mucosa (Right), whereas low BMP4 expression was observed. Yellow circles show nerve cells and the black contour shows RNA-ISH in high magnification. Abbreviations: FB, fibroblasts; SM: smooth muscle; MFB: myofibroblasts.

We compared our transcription profiling of stomach fibroblasts with that of the intestinal fibroblasts in the cross-tissue fibroblast atlas (Buechler et al., 2021). Although PDGFRA^hi^ fibroblasts were limited in intestinal tissues in the public database, our scRNA-seq data of stomach fibroblasts included PDGFR+ fibroblasts (Figure 5B; Figure S4A). These PDGFR+ stomach fibroblasts and the corresponding PDGFRA^hi^ intestinal fibroblasts in the public dataset commonly have specific expression signatures of, for example, *PDGFRA*, *WNT5A,* and *BMP4*. In addition, the numbers of metaplastic enterocytes and PDGFR+ fibroblasts were positively correlated in our single-cell dataset (Figure S4B), suggesting that metaplastic epithelial cell and PDGFR+ fibroblasts interact with each other biologically to maintain the metaplastic intestinal differentiation of the stomach.

To test the aforementioned hypothesis, we conducted RNA-ISH of *BMP4*, one of the signature genes among the PDGFR+ fibroblasts, and *NOG*, an intrinsic BMP antagonist (Zimmerman et al., 1996). In intestinal metaplastic mucosa, *BMP4*+ fibroblasts were more frequently discovered surrounding the metaplastic epithelial cells in the surface areas than in the gastric mucosa (Figure 5C; Figure S4C). The frequent existence of *BMP4*+ fibroblasts was also detected in normal gastric mucosa adjacent to metaplasia (Figure 5C; Figure S4C); however, in normal gastric mucosa distant from IM, *BMP4*+ fibroblasts were found infrequently (Figure 5C; Figure S4C). This suggests that composition changes in the population of specific fibroblasts occur earlier than the epithelial changes over the course of IM. In our ISH analysis, *BMP4* expression levels of the fibroblasts in the metaplastic mucosa of the stomach were comparable with those in genuine colonic mucosa (Figure 5C), suggesting that the physiology in BMP4-related tissue homeostasis was similar in metaplastic stomach glands and colorectal crypts, confirming by 3 stomach specimens including IM gland and 3 colon specimens. Also, we calculated the ratio of the BMP4 green signal area of RNA-ISH in the stroma from randomly selected five fields of the mucosal surface in normal gastric mucosa, transitional mucosa, intestinal metaplastic mucosa, and colonic mucosa. We found the monotonically increasing of BMP4 from normal gastric mucosa to metaplastic and colonic mucosa (p= 0.001445; Figure S4D). With these findings, we showed, for the first time, our hypothesis that the increase of BMP4 in the fibroblasts precede and may even induce IM. *NOG* expression was neither obvious in our scRNA-seq analysis nor was it observed in any cells in the mucosal layers of the stomach (Figure 5C; Figure S4A); however, neuron cell clusters, including ganglion cells, in submucosal layers expressed *NOG* in the stomach (Figure 5D). Drokhlyansky et al. (2020) showed that *NOG* is expressed in neuron cells in the colonic submucosa, but we are the first to report that neuron cells are the intrinsic source of *NOG* in the stomach. A previous study found that enteric neural crest cells promote antral stomach organoids (Eicher et al., 2022), suggesting the importance of nerve cells in the development and/or maintenance of epithelial cells. BMP signaling is known to induce *CDX2* expression in the gastric epithelium (Yoon et al., 2016), and NOG is essential for establishing proper gastric organoids (Eicher et al., 2022; Zhang et al., 2020).

In our dataset, KLF+ fibroblasts (Figure 5A) characteristically expressed *SFRP1, SFRP2, PI16,* and *CD34* (Figure 5B; Figure S4A), but no such fibroblasts existed in the intestine in the public fibroblast atlas, suggesting that the KLF+ fibroblasts in our dataset are unique to the gastric mucosa. RNA-ISH of SFRP1 showed KLF+ fibroblasts existed in the submucosa (Figure S5B). *CD34* is a stemness-associated marker not only in hematopoietic cells but also in other mesenchymal cells (Sydney et al., 2014); therefore, KLF+ gastric fibroblasts might have the potential to develop into other fibroblast clusters. Additionally, KLF+ fibroblasts had the highest stemness score in our analyses (Figure S4E), suggesting that these cells are tissue-resident fibroblasts with stemness features, which supports our hypothesis that they have the potential to differentiate into other subtypes.

FibSmo cells (He et al., 2020), which express both fibroblastic and smooth muscle markers (Figure 5B; Figure S4A), are an uncharacterized fraction. They are distributed in the midpoint between myofibroblasts and other characteristic fibroblasts with signature gene expression, such as that of *PDGFRA* and *KLF4* (Figure 5A), suggesting that FibSmo cells play a unique role among the fibroblast lineages. FibSmo cells specifically express high levels of *HHIP* (hedgehog interaction protein; Figure 5B; Figure S4A), which functions as a regulatory component of the Hedgehog signaling pathway, a pathway required in the development of the normal stomach in both mice and human (El-Zaatari et al., 2009; Katoh and Katoh, 2006). We also found that expression of *BARX1*, which encodes a stomach fibroblast-specific transcription factor, was highest in FibSmo cells (Figure 5B; Figure S4A). Kim et al. (2005) found that *Barx1*-knockout mice did not develop a normal stomach and that intestinal markers were activated; therefore, *BARX1*+ stomach fibroblasts may play a key role in normal gastric development. We performed RNA-ISH to confirm the stomach-specific expression of *BARX1* as well as the coexpression of *HHIP* in *BARX1*+ fibroblasts, finding that *BARX1* expression in fibroblasts occurred in the lamina propria of the pyloric and fundic gland mucosa, indicating that *BARX1* interacted with epithelial cells through its downstream genes (Figure 6A). In contrast, *BARX1*+ fibroblasts were not observed in genuine colonic and small intestinal mucosa (Figure 6A); however, in the intestinal metaplastic mucosa of the stomach, the frequencies of *BARX1*+ fibroblasts were comparable to those detected in the gastric mucosa (Figure 6A). We also found clear coexpression of *BARX1* and *HHIP* in parts of the stomach fibroblasts, although *BARX1* expression was detected more broadly than *HHIP* expression in fibroblasts (Figure 6A). Based on these findings, we confirm the existence of a unique subset of stomach-specific fibroblasts, i.e., *BARX1*+/*HHIP*+ FibSmo fibroblasts, which warrant further functional investigation in relation to the development of the stomach and IM.

**Figure 6.**
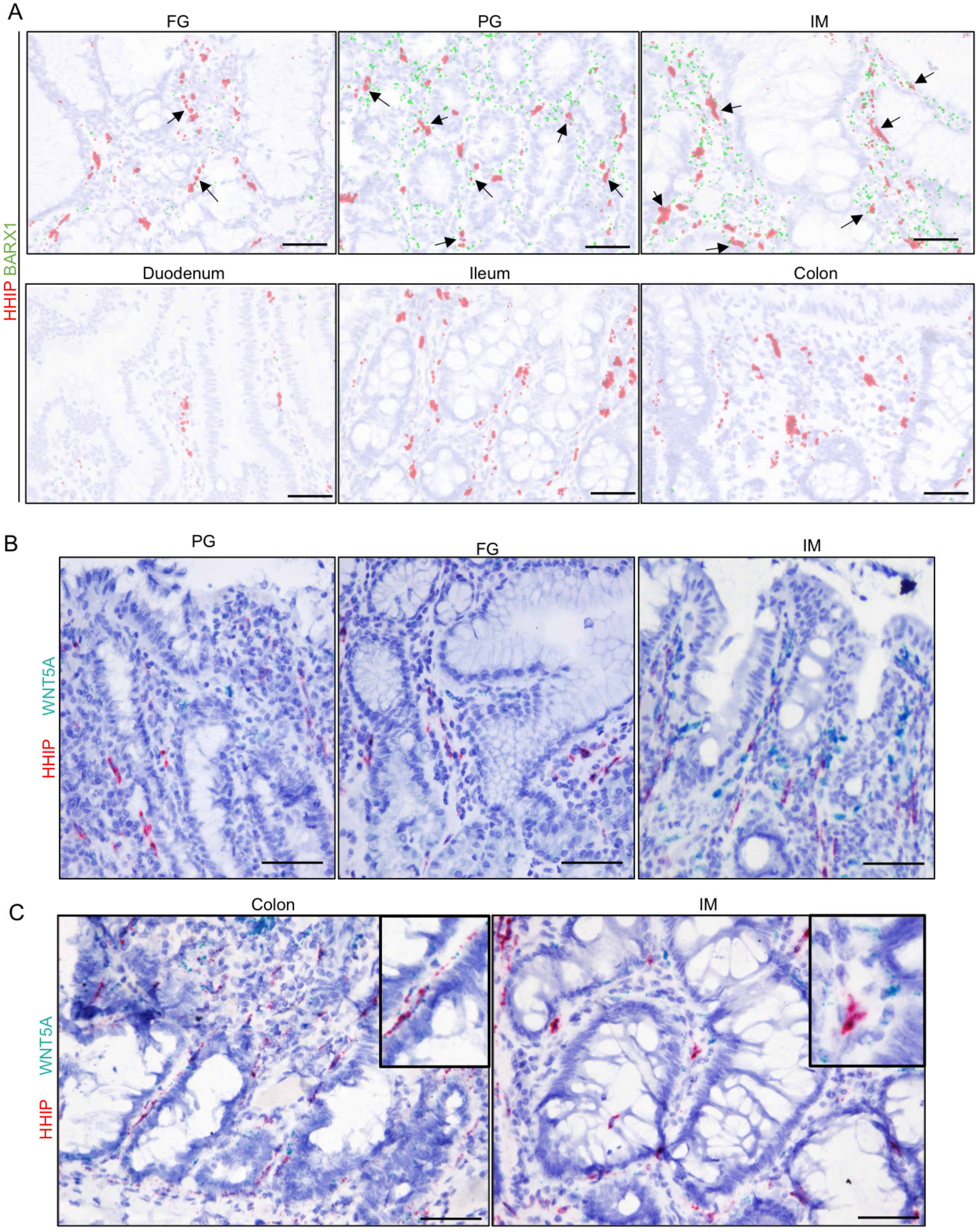
Unique spatial distribution of BARX1, HHIP, and WNT5A-positive fibroblasts in gastric mucosa. (A) HHIP and BARX1 RNA-ISH in each gland or tissue. HHIP and BARX1 coexpression was observed in gastric mucosa including IM (arrows). BARX1 expression was not observed in small intestinal and colonic mucosa, as expected. BARX1 blue signals and HHIP red signals were converted to enlarged green and red pixels, respectively, for ease of detection, as explained in Figure 4D. Arrows: Coexpression of HHIP and BARX1. Scale bar: 50 μm. (B) WNT5A and HHIP RNA-ISH in each tissue. WNT5A was highly expressed in IM relative to its expression in normal mucosa. WNT5A expression was more observed in the surface area and in the just behind epithelial cells, whereas HHIP expression was distant from epithelial cell layers. Scale bar: 50 μm. (C) WNT5A and HHIP RNA-ISH in the colonic mucosa and metaplastic mucosa. WNT5A and HHIP coexpression was observed in colonic mucosa, whereas not in IM. Inset shows higher magnification of each region. Scale bar: 50 μm.

*BARX1* is reported to induce the expression of secreted frizzled-related protein (SFRP), which promotes differentiation of the stomach epithelium by blocking local WNT signaling during the developmental stage (Kim et al., 2005). Interestingly, *BARX1* was expressed at measurable levels in all fibroblast subtypes in our samples, but SFRP expression was mostly limited in the KLF+ fibroblasts (Figure 5B; Figure S4A). RNA-ISH revealed that SFRP1 and BARX1 were not coexpressed in stomach fibroblasts, suggesting that BARX1 does not induce SFRP1 in the fibroblasts of the adult stomach (Figure S5A). Additionally, SFRP+ fibroblasts were observed only in the submucosal layer of the stomach, unlike in the colon mucosa in which SFRP was expressed by fibroblasts in the mucosal layer (Figures S5B and S5C). Therefore, we might postulate the hypothesis that insufficient WNT inhibition by SFRPs in intestinal metaplastic mucosa, unlike in normal colorectal mucosa, could be one of possible causes of unregulated malignant transformation of the metaplastic stomach epithelia.

Our scRNA-seq analysis confirmed that some fractions of stomach fibroblasts are sources of cytokines that regulate WNT or Hedgehog signaling, i.e., the PDGFR+ fibroblasts and FibSmo cells express *WNT5A*, an inhibitor of canonical WNT signaling, and *HHIP*, respectively (Figure 5B). Several studies have suggested that the physical distributions of fibroblasts and resultant local cytokine milieus are vital to the proper development of digestive organs (Ormestad et al., 2006; Roy et al., 2016; Shinohara et al., 2010). SHH or IHH mutant stomachs show abnormal morphology or expression of intestinal markers (Thompson et al., 2018); therefore, the pan-Hedgehog inhibitor HHIP has important functions in the development and homeostasis of the stomach. Thus, we speculate that the spatial distributions of specific stomach fibroblast subtypes play important roles in the tissue architectures of gastric and metaplastic mucosa. Our ISH analysis of *WNT5A* in the stomach and colon tissues revealed that *WNT5A* was expressed in both gastric and metaplastic mucosa, mainly in the surface areas, as reported previously in relation to colonic mucosa (Gregorieff et al., 2005) (Figures 6B and 6C). *WNT5A* expression was more intense in the metaplastic regions than in the pyloric or fundic mucosa, consistent with *BMP4* expression (Figure 6B). *HHIP* expression was also observed broadly in both gastric and metaplastic mucosa; however, in contrast to *WNT5A*, *HHIP*+ fibroblasts were not limited to the surface areas but found at broader depths of the stomach mucosa (Figure 6C). Intriguingly, the close lining of *WNT5A*+ fibroblasts just behind the epithelial cells was frequently observed throughout the stomach mucosa (Figure 6B). Conversely, *HHIP*+ fibroblasts were distributed in stromal spaces at distances from the epithelial cell layers (Figures 6A and 6B). Therefore, WNT5A seems to act locally, i.e., only on the neighboring epithelial cells, whereas HHIP acts more broadly by spreading to distant epithelial cells. Notably, although *HHIP+/WNT5A*+ double-positive fibroblasts were frequently observed in colonic mucosa, such coexpression of *HHIP* and *WNT5A* was not found among stomach fibroblasts, regardless of gastric or metaplastic conditions (Figure 6C). In conclusion, although gastric and colonic fibroblasts share some characteristics, they also harbor their own specific features and probably have distinct functions.

We performed gene regulatory network analysis on the fibroblasts (Figure S5D), finding that forkhead box transcription factors play a role in their biology. Specifically, FOXF1, FOXF2, and FOXL1 transcription activities were upregulated in PDGFR+ fibroblasts as well as FibSmo cells. Correlations among *FOXF1, FOXF2*, and *BMP4* expression and between *FOXL1* and WNT expression have been reported in murine colon fibroblasts (Kaestner, 2019; Ormestad et al., 2006). Therefore, FOXF1/2+ and FOXL1+ fibroblasts are apparently important for regulating stomach homeostasis and metaplastic transformation.

### Immune and endothelial cells

We obtained 71,594 B and plasma cells, including 41,544 plasma cells (with characteristic expression of immunoglobulins, such as *IGHG1, IGHA1,* and *IGKC*), 29,816 B cells (*MS4A1* and *HLA-DRA*), and 15,778 T lymphocytes, with 5 subclusters: CD8 T cells (*CD8A* and *CD8B*; including the GZMK+ CD8 subtype), CD4+CTLA4+T cells (*CD4, PDCD1, CTLA4,* and *FOXP3*), CD4+CD40LG+T cells (*CD4* and *CD40LG*), and γδ T cells (*TRDC* and *TRGC1*) (Figures S6A, S6B, S6E, and S6F). Notably, the proportion of γδ T cells was higher in intestinal metaplastic mucosa compared with that in stomach mucosa (Figure S4B), consistent with the study of Romi et al. (2011), who reported a positive correlation between the number of γδ T cells and *Helicobacter*-associated gastritis. We obtained 2,002 myeloid cells with clusters of dendritic cells, monocytes/neutrophils, and macrophages (Figures S6C and S6G). Conventional M1/2 macrophages or CD14/CD16 monocyte clusters were not identified, consistent with a scRNA-seq atlas of tumor-associated myeloid cells (Cheng et al., 2021), which showed that the myeloid cells have complex phenotypes rather than classical M1 and M2 phenotypes.

In total, 3,028 endothelial cells were identified with 4 subtypes: 1,950 PLVAP+ endothelial cells, 748 ACKR1+ cells, 243 FN1*+* cells (characteristic for *DEPP1, FN1,* and *CXCL2*), and 130 lymphatic endothelial cells (*CCL21* and *LYVE1*) (Figures S6D and S6H). FN1+ cells seemed to be arteries, whereas other endothelial cells were considered venous capillaries based on the expression of EFNB2 and EPHB4 (Kania and Klein, 2016) (Figure S6H). Our IHC analysis showed that ACKR1+ endothelial cells existed in vessels at deeper regions of the stomach mucosa, whereas PLVAP+ endothelial cells constructed vessels in much wider layers of the stomach mucosa, regardless of fundic/pyloric and metaplastic mucosa (Figures S6I–K).

### Cell–cell communication analysis of epithelial cells and stromal cells

Using scRNA-seq analysis, we identified diverse stomach cells in both the physiological and metaplastic mucosa. Various signaling interactions between these cells through cytokines, chemokines, and direct ligand–receptor bindings play important roles in the maintenance of tissue homeostasis in the stomach; thus, we sought to elucidate the cell–cell communication (CCC) networks to deepen our understanding of the cellular diversity of the stomach. CCC analysis was performed by combining a gene expression matrix from our scRNA-seq data with known datasets of ligand–receptor complexes (Jin et al., 2021). Notably, epithelial cell lineages were one of the most enriched cell types from and to which a great diversity of CCC was interconnected (Figure 7A). In addition, we found many more interactions were observed among enterocytes, fibroblasts, and myeloid cells in severe IM, compared with nonsevere IM status (Figures S7A and S7B), suggesting nonepithelial microenvironment contributes development and/or maintenance of epithelial intestinal metaplasia. From the CCC detected around epithelial cells, we focused on LEFTY1+ cells as models for investigating the possible mechanisms by which stem cell properties are maintained (Figure 7A). As shown in Figure S2B, the LEFTY1+ cells also expressed CD44, another known stem cell marker (Ye et al., 2018). CCC analysis revealed that a signaling network of macrophage migration inhibitory factor (MIF), a ligand of the CD74+CD44 complex and CD74+CXCR4 complex (Becker-Herman et al., 2021; Shi et al., 2006), had characteristic features, i.e., many of MIF–(CD74+CD44) and MIF–(CD74+CXCR4) interactions are mediated by LEFTY1+ cells among all the cell types identified in gastric mucosa (Figure 7B). We conclude, therefore, that LEFTY1+ epithelial cells function as a hub of cellular communications via CD44 networks. Given that LEFTY1+ cells also show the highest expression of MIF among epithelial cell types (Figure S8A), CD44, and its communication networks potentially function in an autocrine manner in these cells. The multilayered interactions of LEFTY1+ cells with various other cells, including epithelial cell types, support our hypothesis that LEFTY1+ epithelial cells compose a possible stem cell cluster. CD74 is another communication partner of CD44, and the CD74–CD44 complex functions to prevent apoptosis and maintain stem cell properties (Becker-Herman et al., 2021; Gore et al., 2008; Shi et al., 2006). In contrast to other cell types, we found that a CD44 and CD74 coexpression pattern existed in LEFTY1+ epithelial cells (Figure S8B), providing further evidence that LEFTY1+ possible stem cells function in concert with the CD44 network.

**Figure 7.**
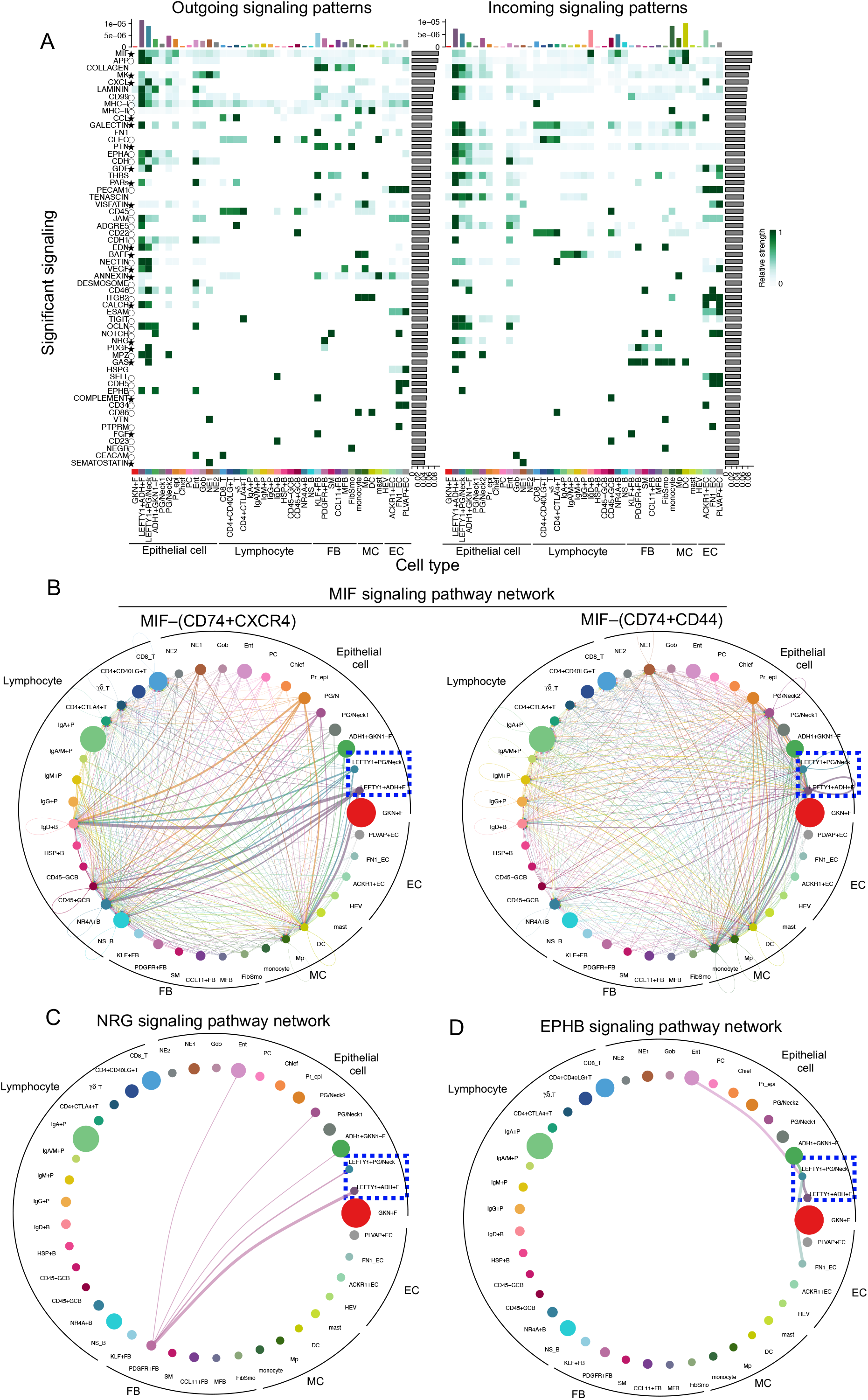
Global view of cell–cell interactions among each cell clusters identified in gastric mucosa, and detailed analysis for focused interactions. (A) Overview of each cell-type and signaling patterns showing that LEFTY1+ cells were centered in the communication network. Top bar plots showing the sum of the communication probability calculated by cellchat library for each cell type. Right bar plots showing the proportion of the contribution in each signal to the total. Heatmaps showing relative strength for each cell type in each signaling. *: secreting signals; O: cell–cell contact; no symbol: extracellular matrix. Jin et al. (2021) provide the signal ligand and receptor pair details. (B–D) Focused interaction network performed by CCC analysis. The circle size of each cluster reflects on the number of cells in each cluster. The thickness of the flow indicates the communication probability. Arrows on outgoing signals and the cluster circles are the same color. (B) Macrophage migration inhibitory factor (MIF) signaling pathway network showing that LEFTY1+ cells expressed both the ligand and receptor. MIF interacts with the CD74+CXCR4 and CD74+CD44 complexes. (C) Neuregulin (NRG) signaling pathway network showing that PDGFR+FB expressed the ligand, NRG1, and enterocytes and that its progenitor cells expressed its receptors ERBB2 and ERBB3. (D) EPHB signaling pathway network showing that LEFTY1+PG/Neck expressed EPHB2 and other mature enterocytes expressed its ligand EFNB2.

As we reported, PDGFR+ fibroblasts expressing BMP4 and/or WNT5A showed characteristic properties in the stomach mucosa, specifically in the intestinal metaplastic mucosa. Additionally, a CCC comparison between severe and nonsevere IM revealed the specific enrichment of NRG signaling in severe IM (Figures S7A and S7B). We also found that PDGFR+ fibroblasts expressed *NRG1*, the product of which is neuregulin-1 (NRG1), characteristically (Figure 7C; Figure S8C). In our CCC analysis of the NRG1 network, PDGFR+ fibroblasts interacted with various epithelial cells, including LEFTY1+PG/Neck cells, LEFTY1+ADH1+F cells, and enterocytes, all of which expressed ERBB3, a receptor for NRG1 (Meyer and Birchmeier, 1995) (Figure S8D). NRG1–ERBB2/ERBB3 signaling is known to act against apoptosis to preserve differentiation in human trophoblasts (Fock et al., 2015), and *NRG1* drives intestinal stem cells to proliferate and regenerate in damaged epithelia (Jardé et al., 2020). Over the course of IM in the damaged stomach, stomach epithelial cells receive NRG1 signals from fibroblasts and eventually differentiate into metaplastic enterocytes. Our CCC analysis is consistent with our spatial distribution analysis of the PDGFR+ fibroblasts (Figures 5C and 6B), i.e., the migration of PDGFR+ stromal fibroblasts precedes the development of intestinal metaplastic mucosa. Taken together, our CCC analyses show that PDGFR+ fibroblasts and secreted NRG1 play important roles in the development of intestinal metaplastic mucosa.

Interactions between EPHB2 in LEFTY1+ PG/Neck cells and EFNB2 in various epithelial and FN1+ endothelial cells were of interest in our CCC analysis (Figure 7D; Figure S8D). Eph–ephrin complexes have a distinct feature of generating bidirectional signals that affect both Eph-expressing and ephrin-expressing cells. Eph–ephrin interactions generate repulsive reactions, which play important roles in the formation of cell clusters and stripe patterns in organogenesis, including in somite and neuronal differentiation (Kania and Klein, 2016; Pitulescu and Adams, 2010). In murine intestinal mucosa, stem cells, during their differentiation into proliferation progenitors, gradually lose *Ephb2* expression, whereas *Efnb2* expression is highest at the villous–crypt boundary (Kania and Klein, 2016). In our scRNA-seq dataset, surface epithelial cells, such as metaplastic enterocytes and ADH1+GKN1-F cells, showed the highest levels of *EFNB2* expression (Figures S8D and S8E). Using IHC, we showed that EPHB2 was only positive in the crypt base of IM, whereas EFNB2 was positive in other regions of IM and the superficial region of gastric mucosa (Figure S8F). This confirms that the function of Eph– ephrin repulsion in intestinal metaplastic mucosa is similar to that in colorectal crypts; however, this interaction was not observed in stomach mucosa. This implies that different combinations of Eph–ephrin interactions or unknown regulation mechanisms might play some roles in stomach glands.

Although our CCC analysis did not identify clear enrichment of the major signaling pathways of BMP or WNT between epithelial cells and fibroblasts, the spatial distribution of WNT5A+ fibroblasts (Figures 6B and 6C) suggested that interesting cellular interactions possibly occur around WNT/BMP. According to RNA-ISH analysis, WNT5A+ fibroblasts were mainly found in the surface area of stomach mucosa (Figures 6B and 6C); however, following closer observations, we found that WNT5A+ fibroblasts were also found in line with LEFTY1+ possible stem cells at the crypt bases of IM (Figure S8G). This close physical interaction between LEFTY1+ cells and WNT5A+ fibroblasts was specific to metaplastic mucosa, i.e., it was not detected in pyloric glands (Figure S8G). LEFTY1 functions as a TGF-β inhibitor (Zabala et al., 2020), whereas WNT5A can potentiate as a TGF-β signaling to control stem cell properties and construct regenerative crypts (Miyoshi et al., 2012); thus, LEFTY1 and WNT5A might work in concert in relation to IM in the stomach. We reported that LEFTY1 IHC revealed two different protein expression patterns; the intense dot pattern indicates the directional secretion of protein into the stromal environment, supporting its biological interaction with the local cytokines/receptors around the stomach gland, such as BMP4 or WNT5A.

### Gene set enrichment analysis of the cells identified via scRNA-seq

To investigate the functional status in each cellular lineage, gene set enrichment analysis was performed for various cell clusters. Some known gene sets identified by Busslinger et al. (2021) with parietal, chief, and gastric neck cell properties were clearly enriched in our parietal, chief, and neck cell populations, respectively, confirming the robustness of our analysis (Figure S9A). Metaplastic enterocytes and LEFTY1+ cells were enriched with gene sets for adipogenesis, fatty acid metabolism, glycolysis, oxidative phosphorylation, and reactive oxygen species pathways. Moreover, a gene set of the MYC pathway “HALLMARK_MYC_TARGETS_V1” was characteristically enriched in LEFTY1+ cells together with multiple gene sets associated with stem cell properties (Figure S9B). In general, quiescent stem cells use glycolysis within a hypoxic niche but context-dependently proliferate and differentiate under other conditions, switching to oxidative phosphorylation (Shyh-Chang and Ng, 2017). The enrichments of metabolic-related gene features and MYC pathway activation suggest that the LEFTY1+ epithelial population, a cluster of a possible stomach stem cells, has a complex and dedicated metabolic regulation pathway; thus, additional studies are required to clarify these complex dynamics.

### Hedgehog signal regulation in gastric mucosa

Hedgehog signaling is an important factor that regulates the development and maintenance of gastrointestinal tracts (Dimmler et al., 2003; Katoh and Katoh, 2006; Thompson et al., 2018), but it was not a focus of the above-reported analysis. A previous study showed that *SHH* mutant mice exhibited intestinal transformation of the stomach (Ramalho-Santos et al., 2000). Additionally, recurrent *MALAT1*-*GLI1* fusion was reported in plexiform fibromyxoma and gastroblastoma, stomach-specific mesenchymal and mixed epithelial–mesenchymal tumors (Graham et al., 2017), suggesting the importance of Hedgehog signaling in the stomach, especially mesenchyme. Both SHH and IHH from epithelial cells are known to activate FOXL1-mediated BMP4 expression in mesenchymal cells (Katoh and Katoh, 2006). In our dataset, SHH expression was limited in gastric lineage, whereas IHH was expressed in both epithelial cells of gastric lineages and metaplastic lineages, respectively (Figure S10A). Although, to our knowledge, studies are lacking on the differential biological functions of *SHH* and *IHH* in the gut, *SHH* and *IHH* may function differently during tissue homeostasis of gastric and metaplastic mucosa. The Hedgehog receptors *PTCH1* and *PTCH2* were expressed occasionally in some PG/Neck1 cells and PDGFR+ fibroblasts, and the downstream effectors of Hedgehog, *GLI1*, *GLI2*, and *GLI3*, were expressed modestly in diverse subtypes of epithelial cells and fibroblasts (Figure S10A). Thus, hedgehog pathways might play important roles in the development and maintenance of stomach tissues. Of the fibroblasts, both FibSmo and PDGFR+ cells expressed *PTCH1*, whereas only FibSmo cells expressed *HHIP*, a negative regulator of the hedgehog pathway (Figures S4A and S10A). *PTCH1* and *HHIP* expression can be induced by a downstream hedgehog signal as a component of negative feedback machinery (Katoh and Katoh, 2006); thus, at least FibSmo fibroblasts seem to exhibit active hedgehog pathways, presumably through the receipt of SHH/IHH from epithelial cells. Considering normal gastric cells and metaplastic enterocytes express *SHH* and *IHH*, respectively, and BMP4 expression is regulated by hedgehog signaling, and BMP4-secreting fibroblasts increase in IM gland, we assume that both SHH and IHH affect the balance between BMP and hedgehog signaling, but the suitable ratio of these components may differ between gastric and intestinal metaplastic mucosa. We hypothesized that, once the balance of hedgehog and BMP signaling in the stomach is disturbed by chronic inflammation or other stimuli, tissues may experience a perturbed cytokine environment that induces IM.

### Upregulation of metabolite-related genes in possible stem cells

In addition to our gene set enrichment analysis, other studies have identified the biological links between stem cell features and the metabolic activity of various pathways (Carey et al., 2015; Cheng et al., 2019; Tischler et al., 2019). Specifically, Cheng et al. (2019) suggested that HMGCS2 and its metabolite β-hydroxybutyrate play important roles in maintaining epithelial stemness, inhibiting histone deacetylase (HDAC), and reinforcing NOTCH signaling in the intestine. They also reported that *HMGCS2* is expressed at higher levels in the *LGR5*+ stem cells of the colon. In our dataset, *HMGCS2* and HDACs (*HDAC1*, *HDAC2*, and *HDAC3*) were clearly enriched in LEFTY1+ cells (Figure S10B), indicating that these cells are possible stomach stem cells, consistent with the findings of the aforementioned studies. Moreover, we found that *IDH1* and *IDH2* were enriched in LEFTY1+ cells (Figure S10C). α-Ketoglutarate is a well-established metabolite of the IDHs, and intracellular α-ketoglutarate is known to maintain the stemness of embryonic stem cells (Carey et al., 2015); thus, the enrichment of IDHs in LEFTY1+ cells is also indicative of their stemness properties.

## CONCLUSION

In this study, we constructed the largest ever stomach cell atlas at a single-cell resolution. This data constitutes a unique resource that will be used in a variety of investigations on stomach development and disease pathology. Combined with sophisticated bioinformatics analyses, we identified a novel candidate common stem cell population in the adult stomach, i.e., the LEFTY1+ cell cluster, and uncovered the skewed and characteristic distributions of specific subtypes of fibroblasts in the ecosystems of normal and metaplastic stomach mucosa. In addition, our CCC analysis revealed that LEFTY1+ is the most enriched population in both outgoing and incoming signaling patterns, supporting the hypothesis that the LEFTY1+ population plays a pivotal role in the maintenance of gastric epithelial integrity. Gene set enrichment analysis and focused metabolic gene expression analyses confirmed and further suggested the complex dynamics of stem cell metabolism. Overall, our study provides novel and unexpected findings related to the normal and metaplastic ecosystems of the stomach that warrant further developmental and cancer research, including the expansion of our dataset with additional scRNA-seq human cell atlases, such as those of other digestive organs and gastric cancers.

## STAR METHODS

Detailed methods are provided in the online version of this paper and include the following:

- KEY RESOURCES TABLE
- RESOURCE AVAILABILITY

- Lead contact
- EXPERIMENTAL MODELS AND SUBJECT DETAILS

- Human specimens
- METHOD DETAILS

- Single-cell RNA sequencing
- Data preprocessing for scRNA-seq
- Clustering, visualization, and cell annotation
- Annotation of IM status
- Transcriptional entropy and trajectory analysis
- Gene regulatory network and cell-cell interaction analysis
- Hematoxylin and eosin staining
- Immunohistochemistry
- RNAScope
- Signal magnification of RNAScope

## ACKNOWLEDGMENTS

We thank Enago (www.enago.jp) for the English language review. Figure illustrations were created in part using <u>Biorender.com</u>. This study was supported by the AMED Practical Research for elucidation of genomic diversity and identification of clinical intervention for gastric cancers with a focus on humoral immunity (grant number JP 21cm0106551 to S.I.). This publication is part of the Human Cell Atlas - www.humancellatlas.org/publications.

## AUTHOR CONTRIBUTIONS

Conceptualization, S.I.; Experiments, A.T., H.K., M.K., A.Y., and T.O.; Computational Analysis, D.K. and A.T.; Resources, H.A., Y.S., and T.U.; Writing, A.T., H.K., D.K., M.K., and S.I.; Supervision, S.I.

## STAR*METHODS

### KEY RESOURCES TABLE

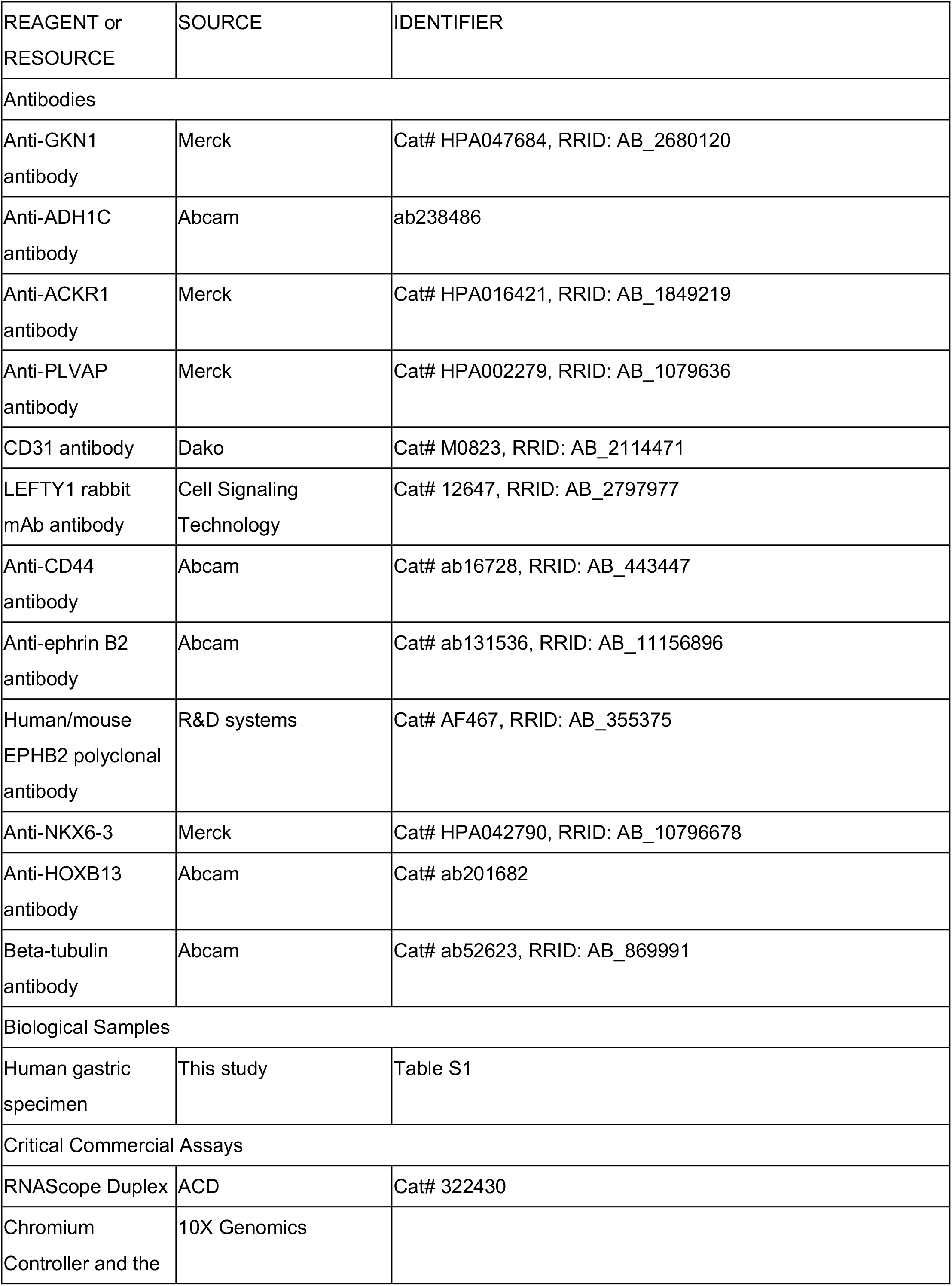
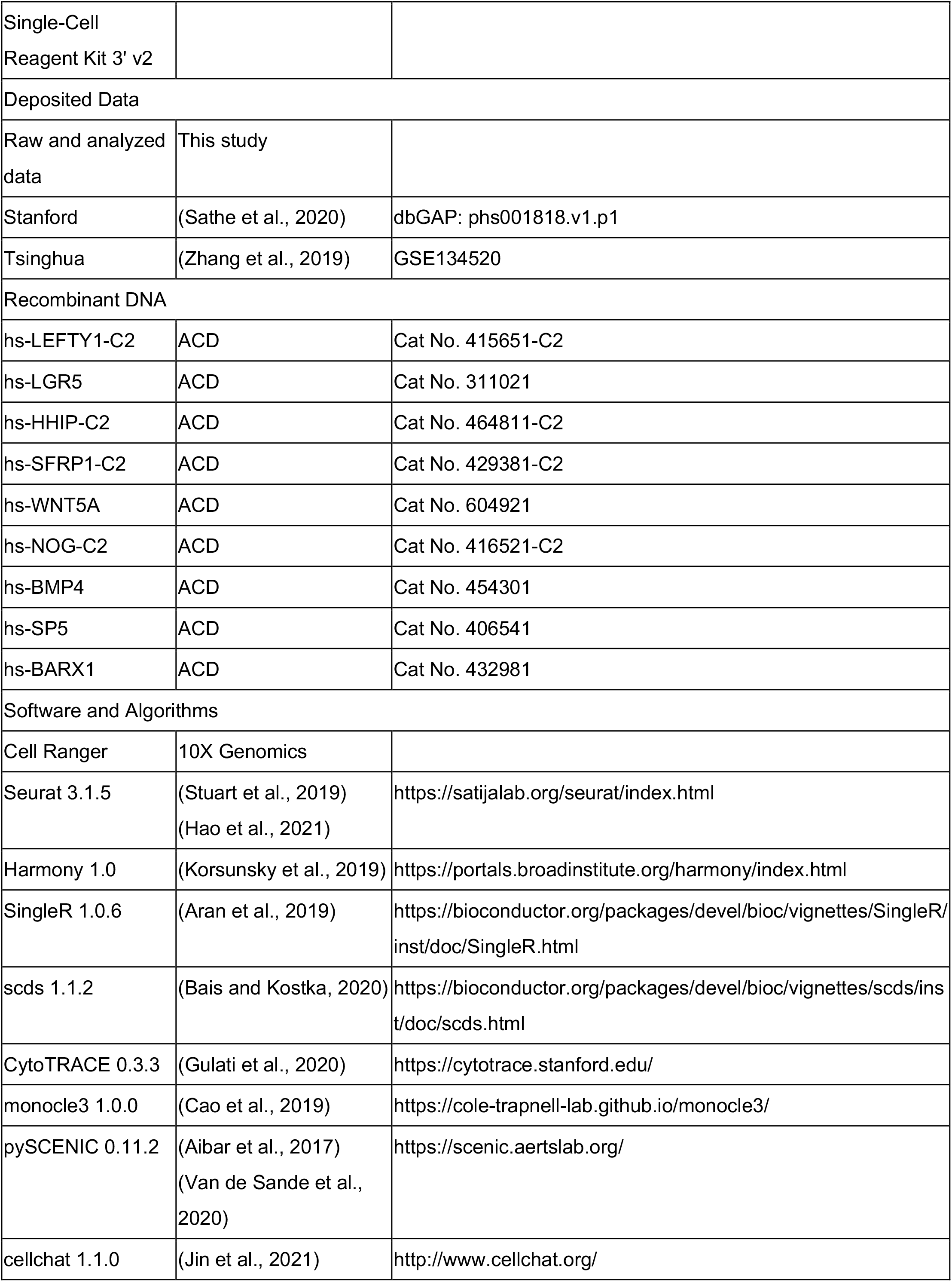

### RESOURCE AVAILABILITY

#### Lead contact

Further information and requests for resources should be directed to and will be fulfilled by the lead contact, Shumpei Ishikawa.

### EXPERIMENTAL MODELS AND SUBJECT DETAILS

#### Materials and methods

##### Human specimens

This study was approved by the Institutional Review Board of The University of Tokyo and written informed consent was obtained from patients. The stomach tissues analyzed in this study were obtained from 15 patients who underwent gastrectomy at The University of Tokyo Hospital from 2017 to 2019. Fresh specimens of noncancerous stomach tissues were obtained immediately after surgeries and subjected to scRNA-seq. Fresh frozen specimens and formalin-fixed and paraffin-embedded (FFPE) specimens of the same patients were also preserved for histopathological examination.

##### Single-cell RNA sequencing

For cases #3–12 and #14–18 listed in Table S1, freshly obtained 3–5-mm-sized specimens from normal stomach tissues were immediately subjected to scRNA-seq. Specimens were cut into pieces of ∼1 mm in size and incubated in collagenase/hyaluronidase (STEMCELL Technologies, Canada) diluted with Dulbecco’s modified Eagle medium [FUJIFILM Wako Pure Chemical Corporation (FUJIFILM), Japan] at 37°C for 30 min with mild agitation, according to the manufacturer’s protocol. Single-cell fractions in the incubated specimens were separated using a 40-μm Cell Strainer (CORNING, USA) with additional filtration conducted using phosphate-buffered saline (PBS; FUJIFILM). The number of cells obtained in single-cell fractions was counted using a hemocytometer (BMS, Japan) following the manufacturer’s protocol. In total, 10,000 cells were subjected to analysis in a Chromium Controller (10X Genomics, USA) following the manufacturer’s instructions. A scRNA-seq library was constructed using Chromium Single-Cell 3′ Reagent Kits ver 2 (10X Genomics), after which the quantification and qualification of sequencing libraries was performed using an Agilent Bioanalyzer (Agilent Technologies, USA) with a High-Sensitivity DNA Kit LabChip (Agilent Technologies).

For cases #7–12 and #14–18, we purified B cell populations used for another experimental purpose (not analyzed in the present study) from the residual cell fractions of the above-mentioned scRNA-seq experiments. B cell fractions were purified using an EasySep Human CD19 Positive Selection Kit II (STEMCELL Technologies) according to the manufacturer’s protocol, after which the cells were resuspended in PBS (FUJIFILM). The purity of the magnet bead-based cellular purifications was considered relatively low; thus, it was considered that a substantial number of mixed cellular populations other than B cells were still included in these samples. Therefore, we included the scRNA-seq data of these B cell-purified samples in combination with the above-mentioned scRNA-seq only when the enriched B cell populations did not affect the purposes and results of our data analyses. Thereafter, 10,000 cells were subjected to scRNA-seq using the Chromium Controller (10X Genomics) following the manufacturer’s instructions. A scRNA-seq library was constructed using Chromium Single-Cell V(D)J Reagent Kits (10X Genomics) combined with Chromium Single-Cell 5′ Library & Gel Bead Kits (10X Genomics), after which the sequencing libraries were quantified and qualified using an Agilent Bioanalyzer (Agilent Technologies) with a High-Sensitivity DNA Kit LabChip (Agilent Technologies).

The entire scRNA-seq library was subjected to next-generation sequencing using an Illumina NovaSeq platform (Illumina, USA). This sequencing was conducted by iLAC (Ibaraki, Japan).

##### Data preprocessing for scRNA-seq

Sequencing data were aligned to human genome GRCh38, and the unique molecular identifiers for each cell were counted using Cell Ranger version 3.1 (10X Genomics). Ambient RNA removal was performed with SoupX version 1.3.7 (Young and Behjati, 2020) using hemoglobin genes and immunoglobulin genes to estimate contamination fractions. Gene expression matrices of Stanford University and Tsinghua University were downloaded from https://www.ncbi.nlm.nih.gov/projects/gap/cgi-bin/study.cgi?study_id=phs001818.v1.p1 and https://www.ncbi.nlm.nih.gov/geo/query/acc.cgi?acc=GSE134520, respectively. All data were then merged using Seurat R package version 3.1.5 (Stuart et al., 2019), and the batch effects among the samples were removed using Harmony R package version 1.0 (Korsunsky et al., 2019). Cells with low quality were then filtered out based on the proportion of mitochondrial gene counts. We used a cell-type specific cutoff value based on the following procedure. First, we inferred the cell types using SingleR version 1.0.6 (Aran et al., 2019) and reference transcriptomic datasets. We then removed cells with >25% mitochondrial genes in the epithelial cell lineage and >15% mitochondrial genes in other cells. Finally, we used scds R package version 1.1.2 (Bais and Kostka, 2020) for doublet cell detection.

We applied the “LogNormalize” function that normalized the gene expression of each cell according to the total expression, multiplied this by a scale factor 10,000, and log-transformed the result using the NormalizeData() function in Seurat.

##### Clustering, visualization, and cell annotation

Principal components were calculated based on the normalized gene expression profiles. The number of principal components was determined using a Jackstraw plot, the p-value thresholds of which were 0.05. tSNE and UMAP dimensionality reduction was performed using the Seurat functions “RunTSNE” and “RunUMAP,” respectively. Cell clusters were identified using the “FindClusters” function in Seurat. Differentially expressed genes were obtained using the “FindAllMarkers” function via MAST (Finak et al., 2015) with the number of genes detected in each cell used as a latent variable. The cell cycle phases for each cell were estimated based on the gene expression of cell cycle marker genes (Nestorowa et al., 2016).

##### Annotation of IM status

In our dataset, IM status was determined based on histology by experienced pathologists. For the Tsinghua University dataset, because IM status was already annotated by the authors, we used the existing annotations. For the Stanford University dataset, we calculated the percentage of intestinal cells in all epithelial cells for each sample, and the IM status was estimated by comparing this percentage with those of the Tsinghua University dataset and our own dataset.

##### Transcriptional entropy and trajectory analysis

Using the count data of scRNA-seq and cell-type annotations from Seurat as input, CytoTRACE (Gulati et al., 2020) analysis was conducted to calculate transcriptional entropy. Trajectory analysis was performed using monocle3 (Cao et al., 2019) with the same input as that used in CytoTRACE. In monocle3, batch effect removal was conducted using a function implemented in the software (Haghverdi et al., 2018).

##### Gene regulatory network and cell–cell communication analysis

The gene regulatory networks for epithelial and fibroblast cells were inferred using the SCENIC and pySCENIC pipelines (Aibar et al., 2017; Van de Sande et al., 2020). To confirm reproducibility, gene regulatory network analysis was performed 10 times. The Seurat object with cell-type annotation data was converted to a loom object using the loomR library to generate input data for SCENIC. CCC analysis was conducted using Cellchat (Jin et al., 2021) with count data and cell-type annotations used as input data.

##### Hematoxylin and eosin staining

Fresh frozen and FFPE specimens were sliced to thicknesses of 4 μm and subjected to hematoxylin and eosin (H&E) staining. Histopathological slides of fresh frozen specimens were snap-fixed with 4% paraformaldehyde phosphate buffer solution (FUJIFILM) for 10 min at room temperature, and the slides of FFPE specimens were deparaffinized and rehydrated via immersions in xylene (#241-00091; FUJIFILM) and ethanol (#057-00451; FUJIFILM), respectively. Hematoxylin (#6187-4P; Sakura Finetek Japan, Japan) and eosin (#8660; Sakura Finetek) solutions were then used to achieve H&E staining following the manufacturer’s protocols. The stained slides were dehydrated via immersions in ethanol and xylene, respectively, after which glass coverslips (Matsunami Glass, Japan) with Marinol (#4197193, Muto Pure Chemicals, Japan) were used to cover the stained slides. H&E-stained images were then captured using a Hamamatsu Nanozoomer 2.0 HT whole slide scanner (Hamamatsu Photonics K.K., Japan).

##### Immunohistochemistry

For IHC, histopathological slides with FFPE specimens were deparaffinized and rehydrated via immersions in xylene (#241-00091; FUJIFILM) and ethanol (#057-00451; FUJIFILM), respectively. The slides were then autoclaved for 5 min at 120°C while immersed in citrate buffer (pH 6.0) (Abcam, Cambridge, UK). Endogenous peroxidase activity was consumed using 0.3% hydrogen peroxide (Sigma Aldrich, USA) in methanol (#137-01823; FUJIFILM) for 15 min, after which the slides were washed using distilled water. Nonspecific protein–protein reactions were blocked by incubating the slides in 2% bovine serum albumin (Sigma Aldrich, USA)/PBS (FUJIFILM) for 15 min at room temperature. The following primary antibodies were used, which were incubated on the slide at 4°C overnight: GKN1 (1:1,000; Merck, HPA047684), ADH1C (1:2,000; Abcam, ab238486), ACKR1 (1:200; Merck, HPA016421), PLVAP (1:100; Merck, HPA002279), CD31 (1:100; Dako, M0823), LEFTY1 (1:1,000; Cell Signaling Technology, #12647), CD44 (1:200; Abcam, ab157107), EFNB2 (1:200; Abcam, ab131536), EPHB2 (1:200; R&D systems, AF467), NKX6-3 (1:1,000; Merck, HPA042790), HOXB13 (1:3,000; Abcam, ab201682), and TUBB3 (1:200; Abcam, ab52623). After washing the slides with PBS (FUJIFILM) three times, Histostar (Ms+Rb) for Human Tissue (MBL, Japan) was used as a secondary antibody, and the slides were washed using PBS (FUJIFILM) three times. IHC signals were developed using Histostar DAB Substrate Solution (MBL) according to the manufacturer’s protocol. Nuclear staining was performed using a hematoxylin (#6187-4P; Sakura Finetek Japan) solution. The stained slides were dehydrated using immersions in ethanol followed by xylene, after which glass coverslips (Matsunami Glass) with Marinol (#4197193; Muto Pure Chemicals) were used to cover the stained slides. IHC images were captured using a Hamamatsu Nanozoomer 2.0 HT whole slide scanner (Hamamatsu Photonics K.K., Japan).

##### RNAScope

To achieve ISH, a RNAScope 2.5 HD Duplex Assay (Advanced Cell Diagnostics, Inc., Hayward, CA, USA) was performed according to the manufacturer’s instructions. The method used to prepare FFPE samples was the same as that used for IHC analysis. A list of the probes used is provided in the key resources table.

##### Signal magnification of RNAScope

We extracted green or red signals from the image of RNAScope and magnified these signals 100 times using python library cv2 and PIL. Green and red signal thresholds were defined manually.

---

**Table S1.** Sample metadata.

**Figure S1.**
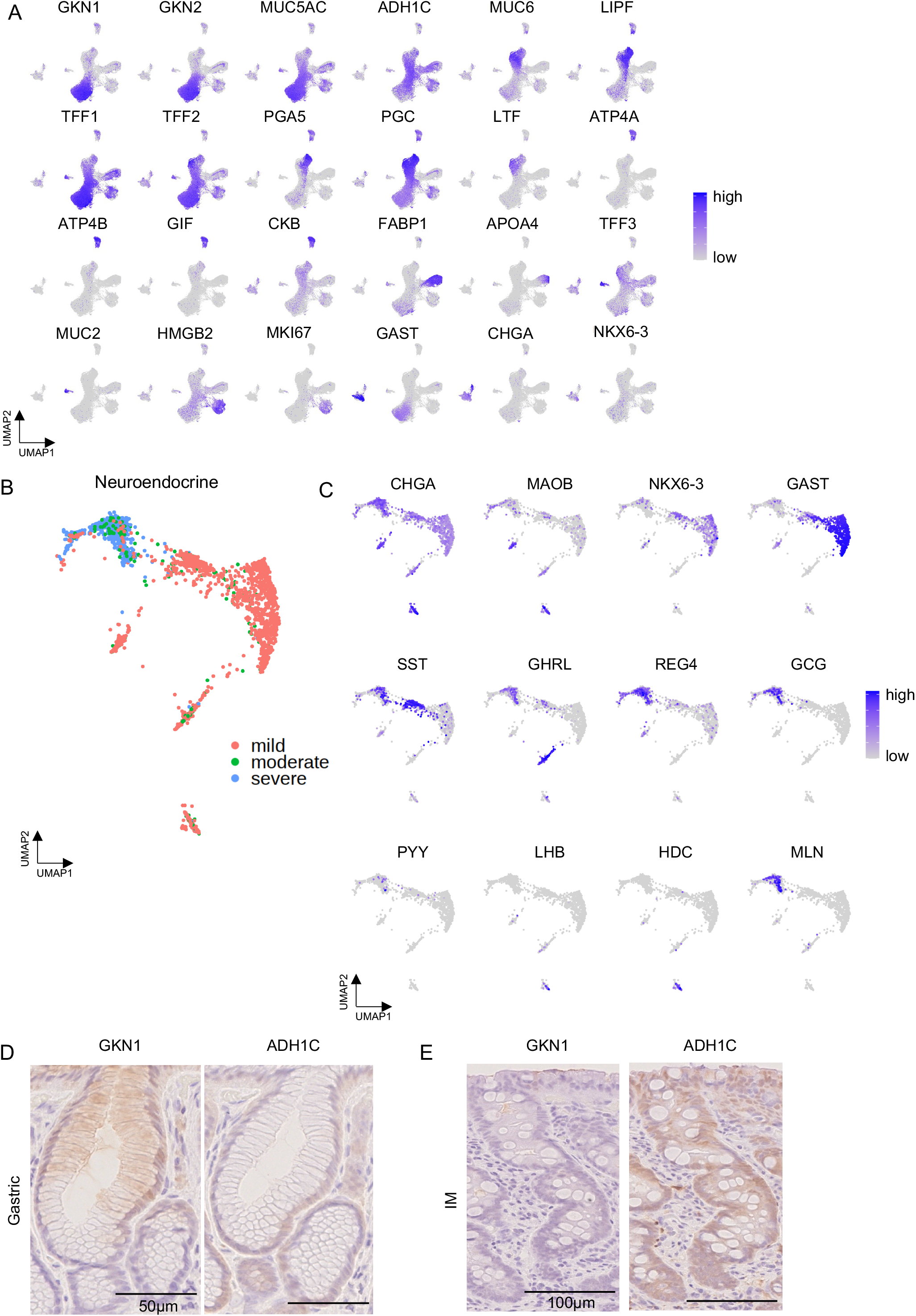
Marker genes of epithelial cells and neuroendocrine-specific cells (related to Figure 1). (A) Representative epithelial cell marker gene expression selected from differentially expressed genes. (B) UMAP of neuroendocrine cells with IM severeness. Almost all neuroendocrine cells from severe or moderate IM samples express REG4. (C) Several enzymes and marker genes of neuroendocrine cells. Almost all *GAST*+ cells are from mild IM samples, and *GCG* and *MLN+* cells are from severe or moderate samples. Rare neuroendocrine cells such as *LHB*+ or *HDC*+ cells were identified. (D, E) IHC of GKN1 and ADH1C in the stomach and IM, respectively. GKN1 expression is limited in the superficial region in normal gastric mucosa. ADH1C expression is in the deeper region in normal gastric mucosa and in all IM region.

**Figure S2.**
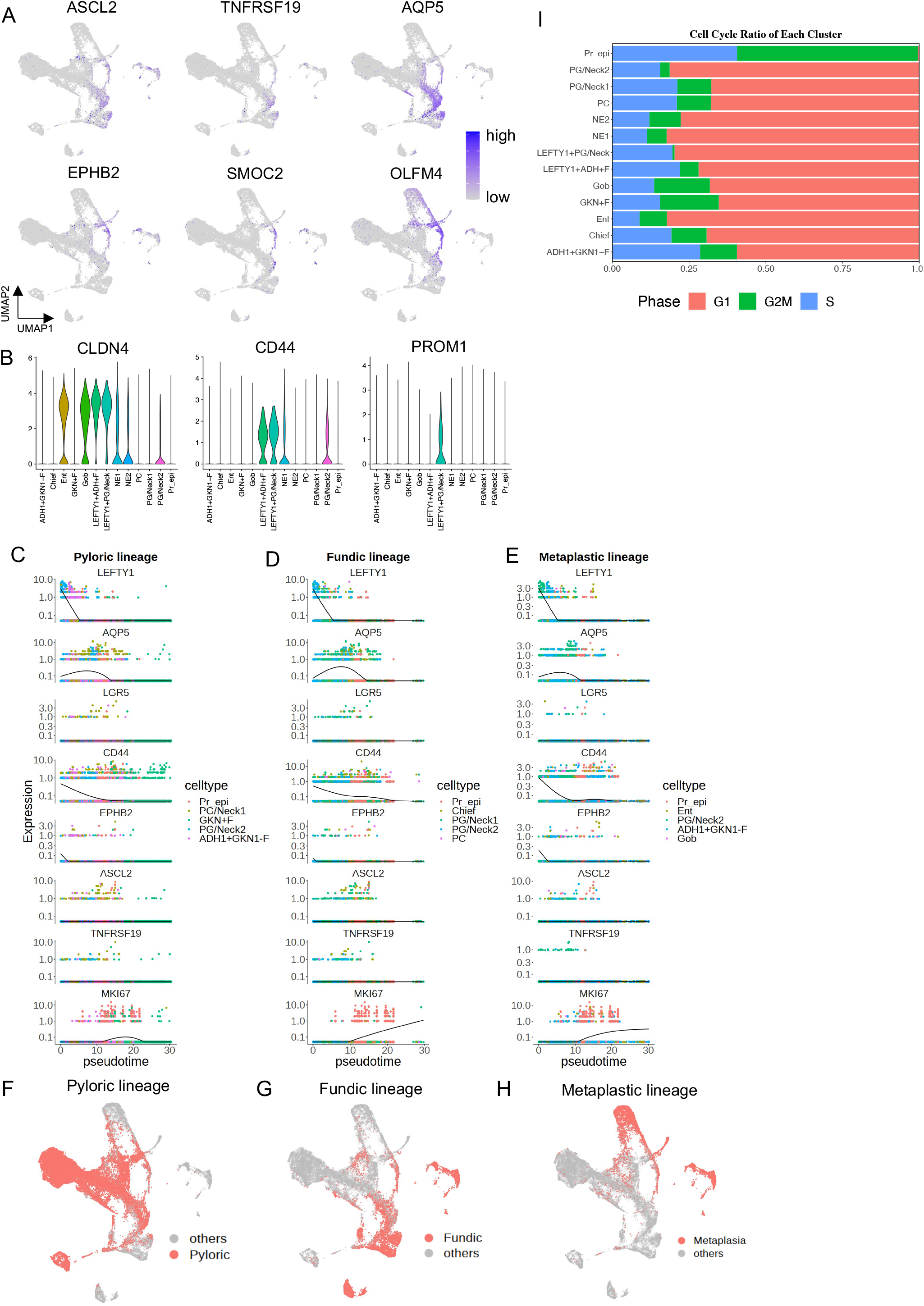
Gastrointestinal stemness-associated genes enriched in LEFTY1+ and pseudotime dynamics (related to Figure 2). (A, B) Stem cell-associated gene expression in the epithelial cells. These marker genes are expressed in LEFTY1+ cells. (C–E) Pseudotime plots of representative stem cell markers in each lineage. LEFTY1 expression is highest in the earliest time in pseudotime among other stemness-associated genes. MKI67 expression is high in the middle of the pseudotime. (F–H) UMAP showing each lineage cell; Pyloric lineages cell include GKN+F, ADH1+GKN1-F, PG/Neck1, PG/Neck2, and Pr_epi; Fundic lineage cells: PG/Neck1, PG/Neck2, PC, Chief, Pr_epi; Metaplastic lineage cells; PG/Neck2, ADH1+GKN1-F, Gob, Ent, Pr_epi. (I) Proportion of cell cycle phases in each cluster. Almost all proliferating epithelial cells show G2M or S phase. The G2M phase was detected less frequently in LEFTY1+ PG/Neck cells, which was compatible with being quiescent. See methods for cell cycle analysis details.

**Figure S3.**
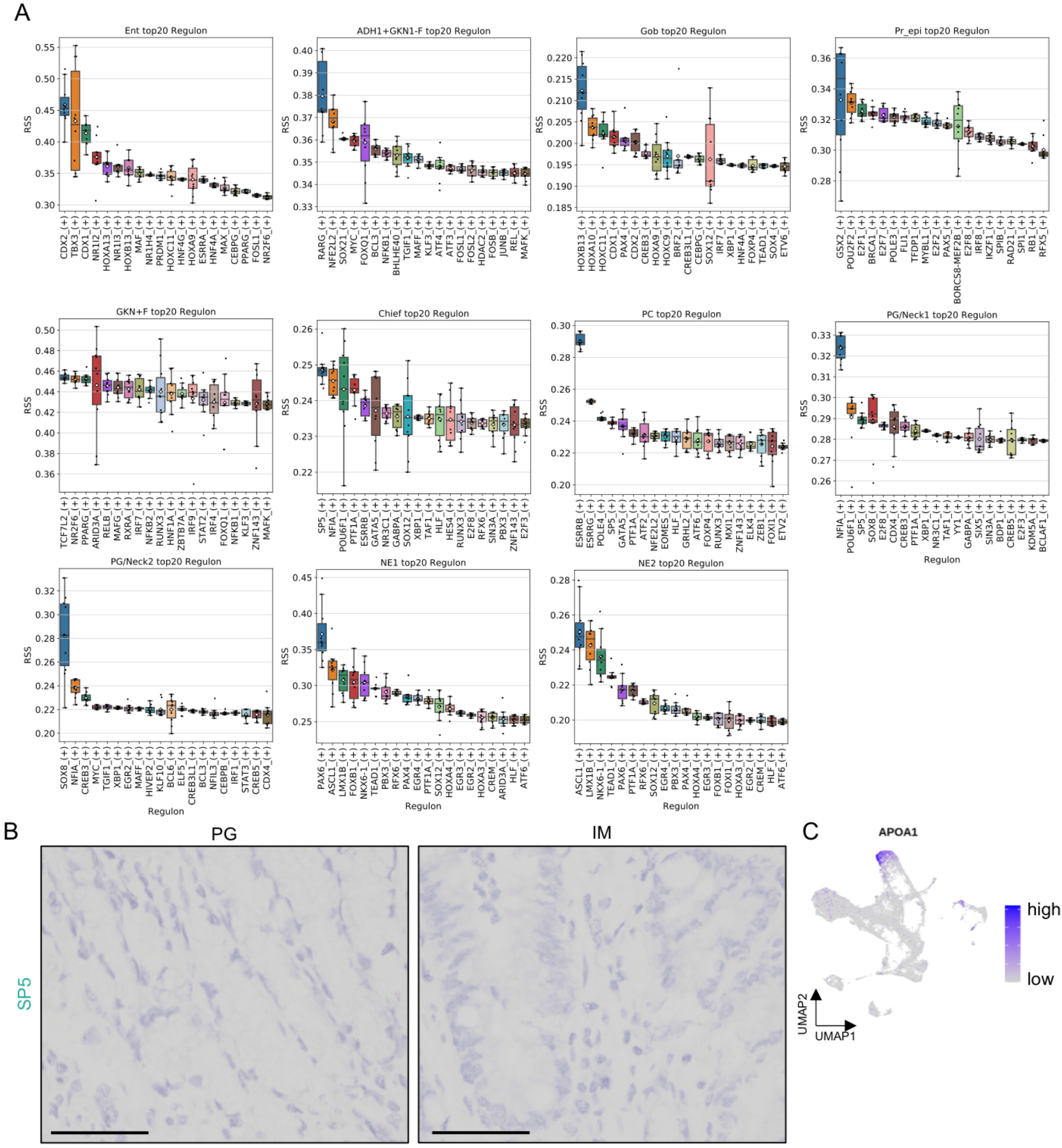
Top 20 regulons, except LEFTY1+ cell clusters, in the epithelial cells (related to Figure 4). (A) Boxplots show the results of gene regulatory network analysis in each cluster for the top 20 regulons with 10 times runs. NE clusters show high activities of ASCL1 and PAX6, PC cluster shows high activities of ESRRB and ESRRG, and Ent cluster shows high activities of CDX2 and CDX1. Fundic gland-specific clusters (PG/Neck1, PC, and chief cells) show a high activity of SP5. (B) RNA-ISH of SP5 showing no signal detection in PG and IM. Scale bar: 50 μm. (C) Expression of APOA1 is limited in the enterocytes.

**Figure S4.**
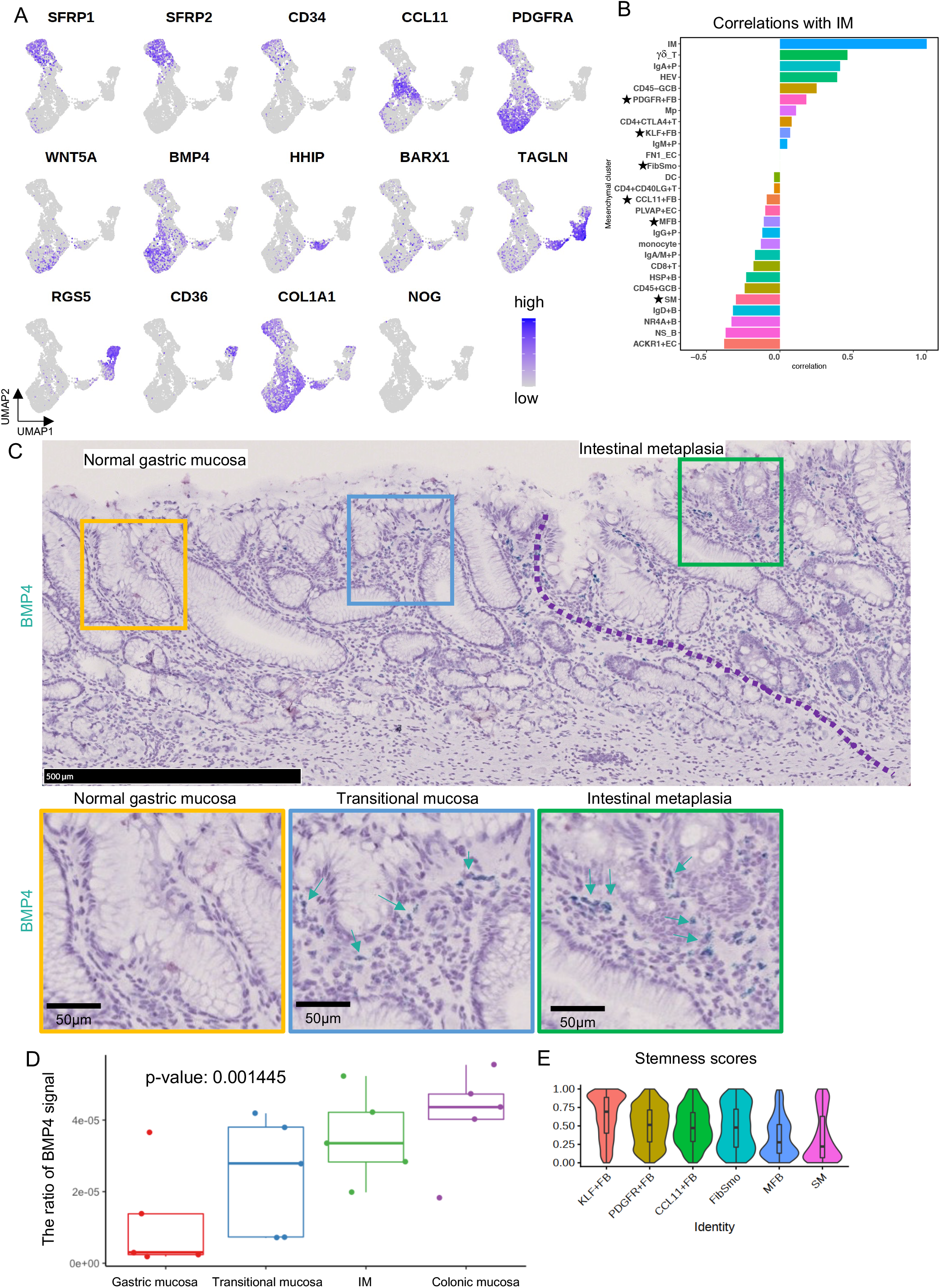
Characteristics of fibroblast subtypes and BMP4-related analysis (related to Figure 5). (A) Representative marker gene expression in fibroblasts. WNT5A and BMP4 expressions are enriched in PDGFR+FB population, and HHIP expression are in FibSmos. (B) Bar plot showing the correlation between the number of metaplastic cells (the total of enterocytes and goblet cells) and ratio of each stromal cell to the number of major clusters to which it belongs. The ratio of PDGFR+FB is positively correlated with the number of metaplastic cells. Each stromal cell ratio was calculated as the number of each subcluster divided by total cell number of the same class. Note: the γδ T cell and PDGFR+FB ratio was calculated as the number of γδ T cells divided by the total number of T cells and the number of PDGFR+FBs divided by the total number of fibroblasts, respectively. *: Fibroblasts (C) Another field of RNA-ISH of BMP4 in IM, transitional mucosa, and gastric mucosa in addition to Figure 4C. In this field of view, the increase of BMP4 expression was also observed in the metaplastic gland and normal gastric mucosa adjacent to IM. Top panel: A low magnification of stomach gland including IM, transitional mucosa, and gastric mucosa. Bottom panel: A high magnification of each gland. The contour colors correspond with the colors of squares in the top panel. Arrows: BMP4 signals. (D) The ratio of BMP4 green signal area in the stromal area in the field of view of the mucosal surface in each gland. The monotonically increase of BMP4 from normal gastric mucosa to metaplastic and colonic mucosa was observed (p=0.001445, two-sided Jonckheere-Terpstra trend test). (E) Stemness scores in the fibroblasts. KLF+FB shows highest score, whereas MFB and SM show lowest scores.

**Figure S5.**
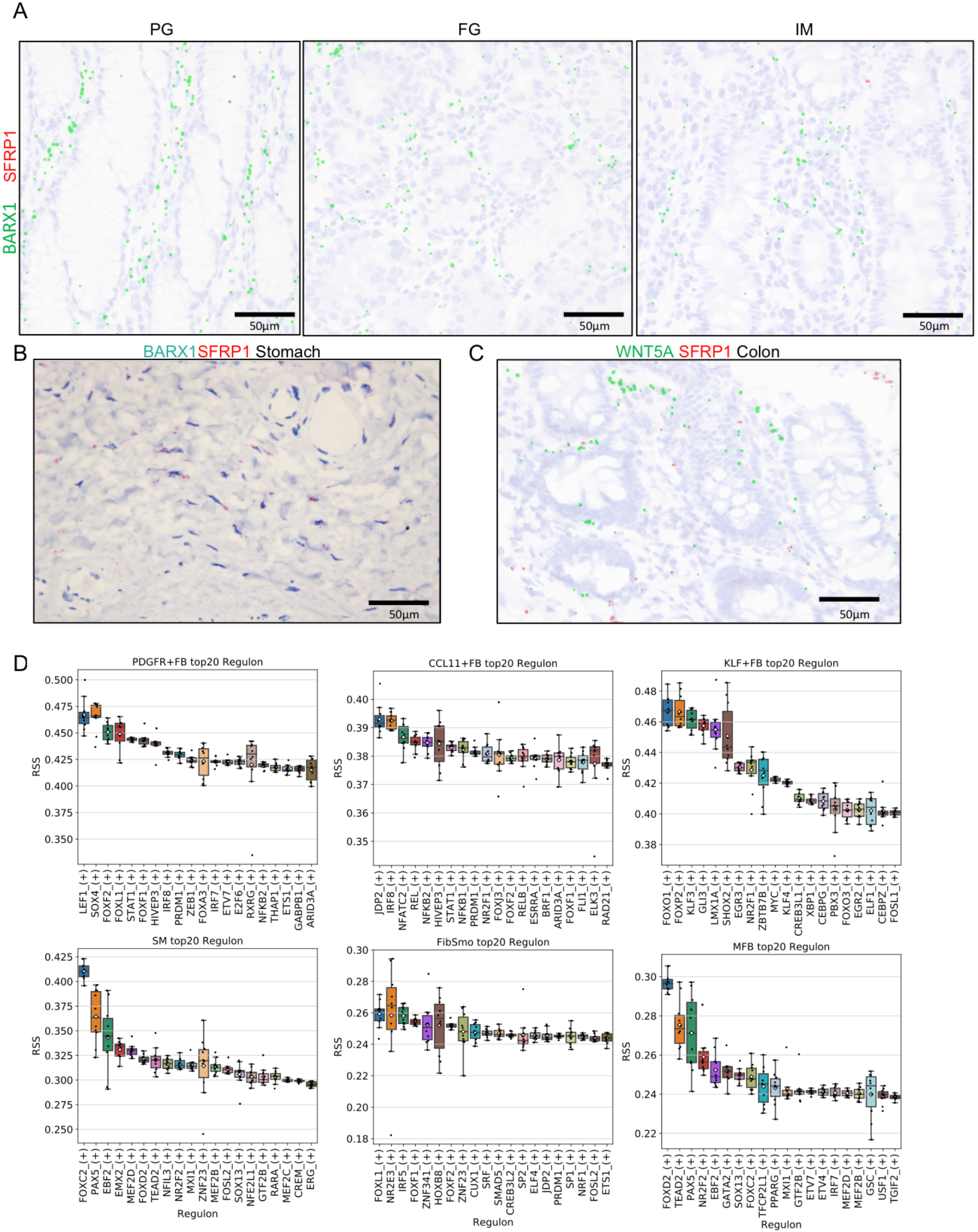
Spatial distribution of BARX1, SFRP1 and WNT5A in the gastric and colonic mucosa and regulon activity of fibroblasts (related to Figure 6). (A) RNA-ISH of SFRP1 (red) and BARX1 (green) showing the lack of SFRP1 and BARX1 coexpression. SFRP1 expression was not observed in the mucosal lamina propria. Each signal was expanded computationally. (B) RNA-ISH of BARX1 (green) and SFRP1 (red) in the gastric submucosal region. SFRP1 expression was limited in the submucosa in gastric mucosa. KLF+FBs existed in the submucosal region because KLF+FBs specifically express SFRP1. (C) RNA-ISH of WNT5A (green) and SFRP1 (red) in colonic mucosa. SFRP1 was expressed in mucosal laminar propria in colonic mucosa but not in stomach mucosa. (D) Boxplots showing the results of gene regulatory network analysis in each cluster for the top 20 regulons with 10 times runs. FOXF1, FOXF2, and FOXL1 transcription activities were upregulated in PDGFR+ fibroblasts as well as FibSmo cells.

**Figure S6.**
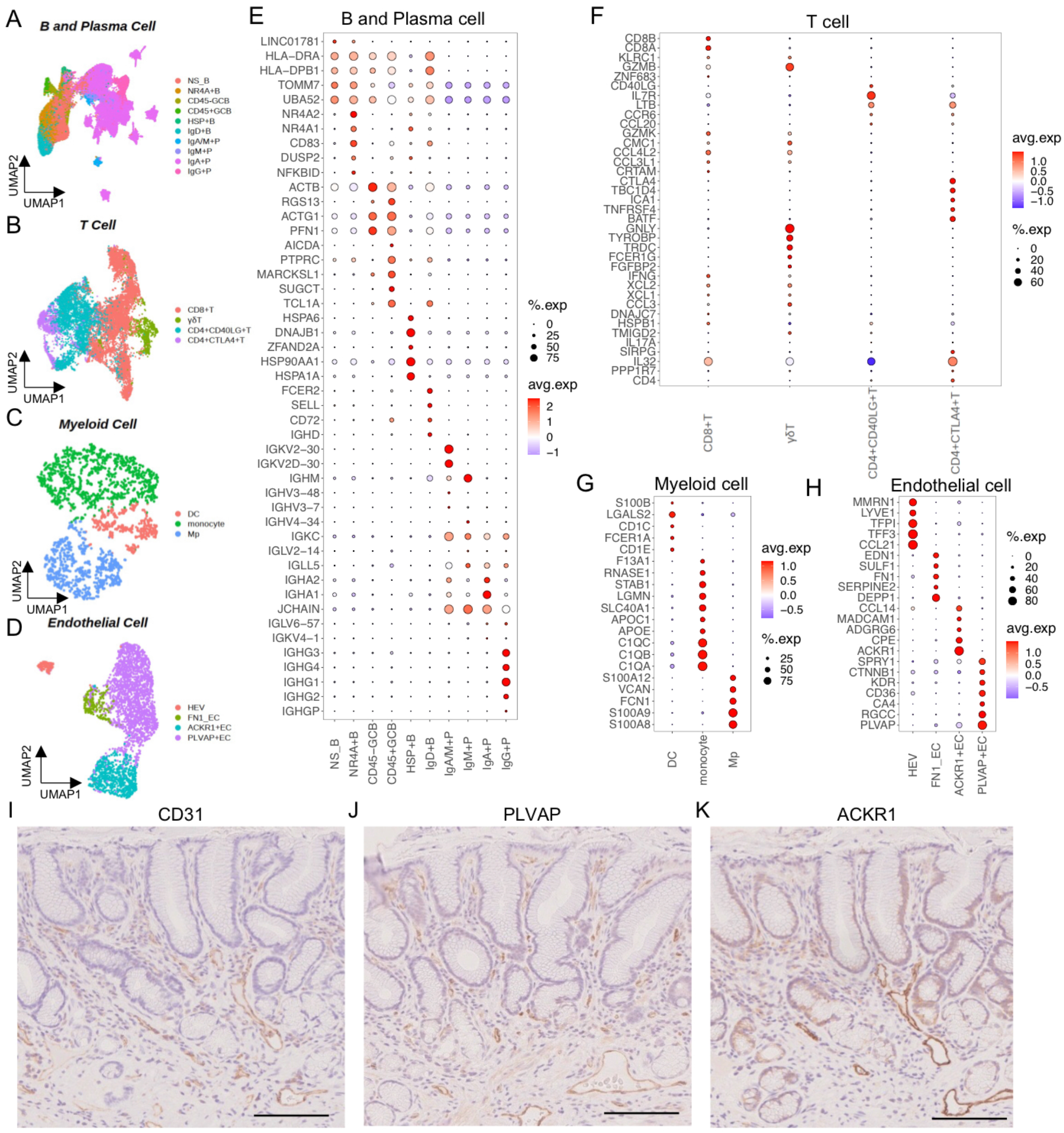
Immune cell and endothelial cell characterization. (A, E) B and plasma cell UMAP and dot plot showing subcluster marker genes. (B, F) T cell UMAP and dot plot showing subcluster marker genes. (C, G) Myeloid cell UMAP and dot plot showing subcluster marker genes. (D, H) Endothelial cell UMAP and dot plot showing subcluster marker genes. (I–K) Images of IHC showing the endothelial marker proteins CD31, PLVAP, and ACKR1. PLVAP was distributed in the entire mucosa. Compared with PLVAP, ACKR1 was found in a deeper region. Scale bar: 100 μm.

**Figure S7.**
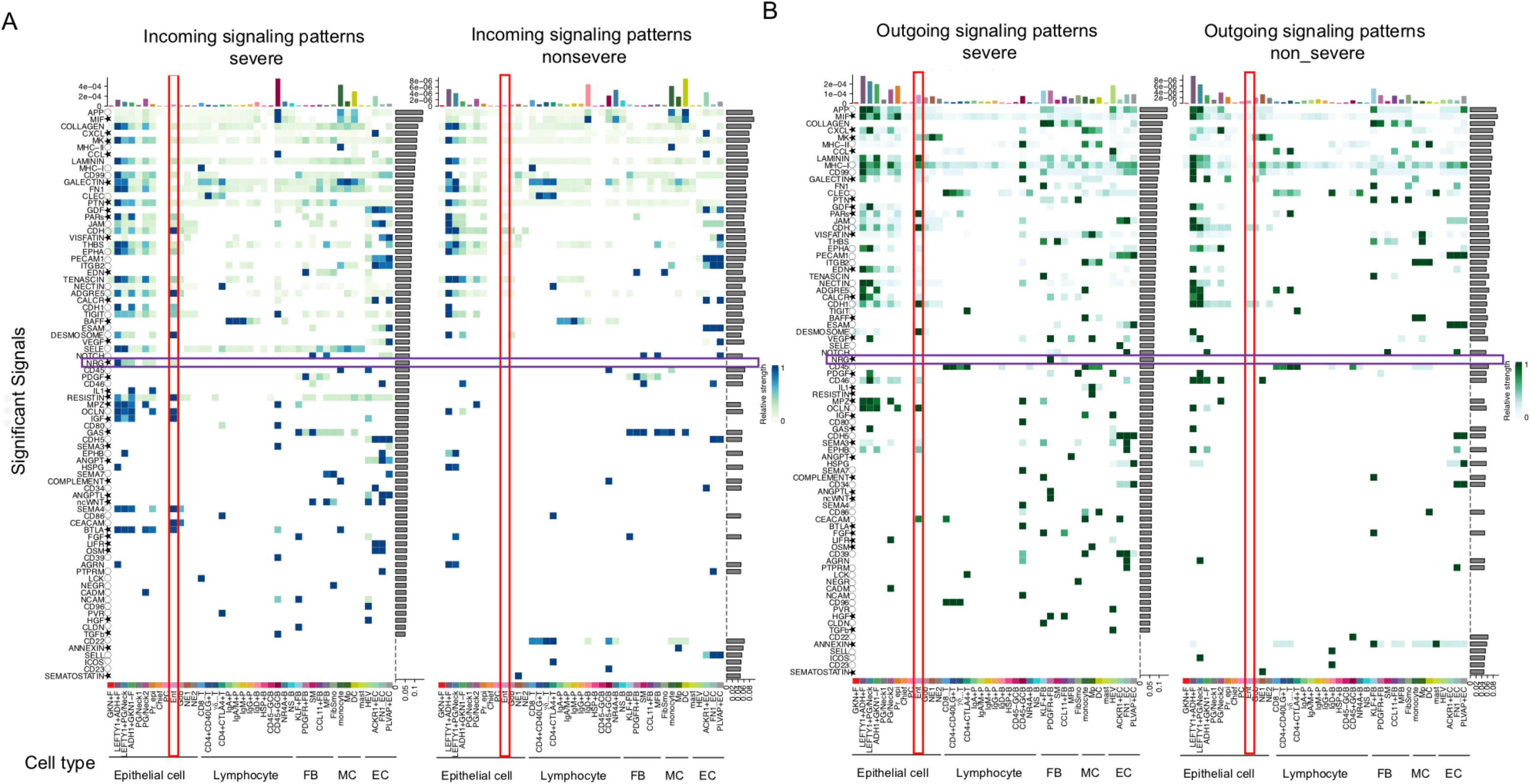
Overview of cell–cell communications between IM severe samples and IM mild/moderate samples (related to Figure 7). (A, B) Overview of each cell-type and signaling patterns showing more interactions in the enterocytes, fibroblasts, and myeloid cells of IM severe samples. Top bar plots showing the sum of the communication probability calculated by cellchat library for each cell type. Right bar plots showing the proportion of the contribution in each signal to the total. Heatmaps showing relative strength for each cell type in each signaling. Red square: enterocytes; purple square: NRG signaling; *: secreting signals; O: cell–cell contact; no symbols: extracellular matrix. Jin et al. (2021) reported the signal ligand and receptor pair details.

**Figure S8.**
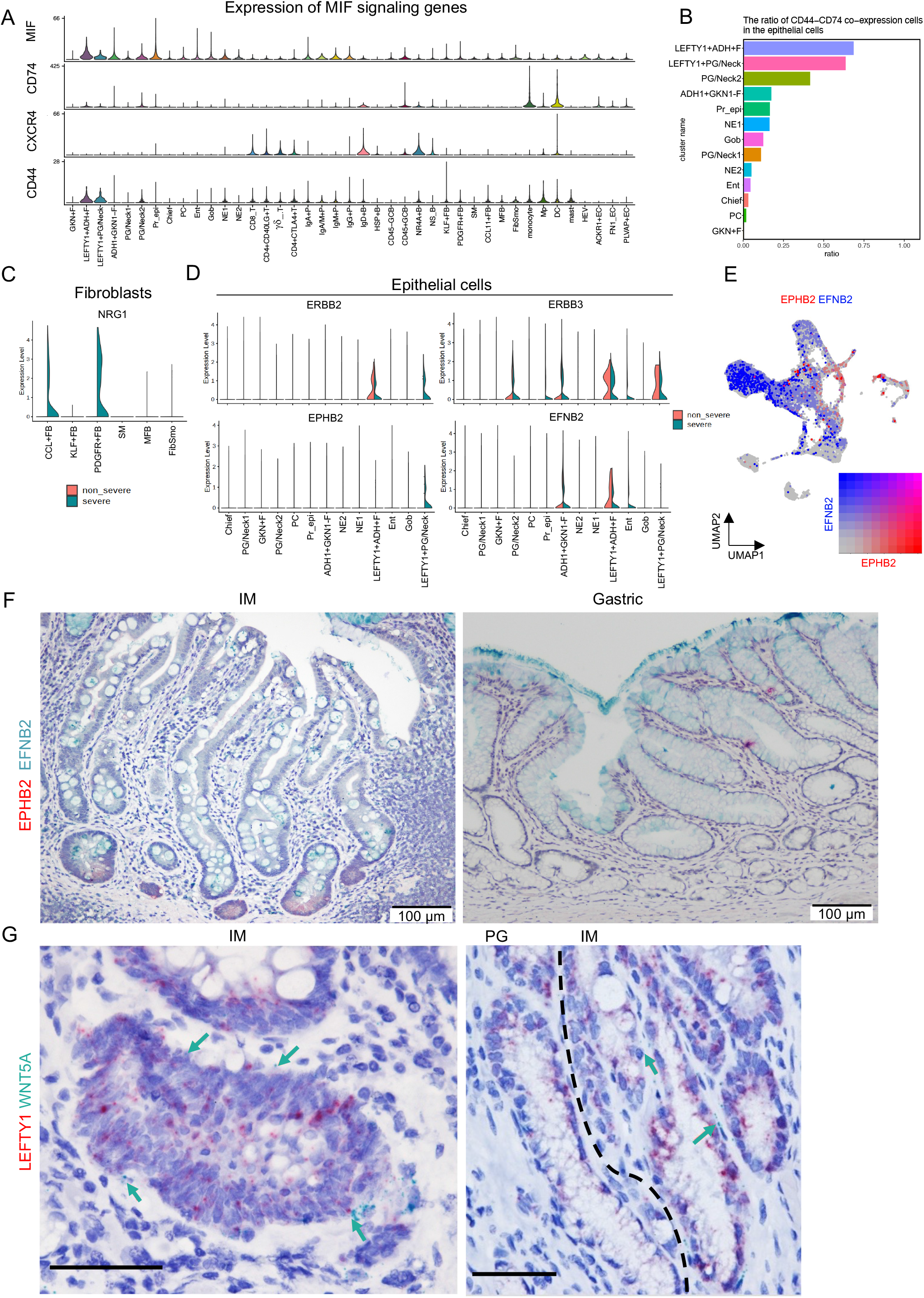
Representative expression of ligand and receptor genes and proteins in the transcriptional profile and human tissue (related to Figure 7). (A) Violin plots showing MIF signaling genes in the all-cluster cells. CD44 and MIF expressions are highest in LEFTY1+cells, whereas CXCR4 expression is highest in B cell clusters. (B) Percentage of cells with CD44–CD74 coexpression in epithelial cells. LEFTY1+cells show higher scores than other epithelial cells. (C) NRG1 expression is limited in PDGFR+FB and CCL11+FB from IM severe samples. Left half: expression of cells derived from IM severe samples; right half: expression of cells derived from IM mild and moderate samples. (D) NRG receptor (ERBB2 and ERBB3) and EPHB signaling (EPHB2 and EFNB2) genes expression in epithelial cells. Left half: expression of cells derived from IM severe samples; right half: expression of cells derived from IM mild and moderate samples. (E) EFNB2 and EPHB2 expression in epithelial cells (blue: EFNB2; red: EPHB2). EPHB2 expression was enriched in LEFTY1+ cells, and EFNB2 expression is higher in GKN1+F and enterocytes. (F) IHC of EFNB2 (green) and EPHB2 (red) in IM and PG. EPHB2 expression was observed in the base crypt of IM samples, and EFNB2 expression was observed in the superficial mucosa both in IM and gastric mucosa. (G) RNA *in situ* hybridization of LEFTY1 (red) and WNT5A (green) in IM and PG. WNT5A expression was observed in the invaginations of IM, showed in Miyoshi et al., 2012. Black dashed line shows the border of IM and the PG. Arrows: WNT5A. Scale bar: 50 μm.

**Figure S9.**
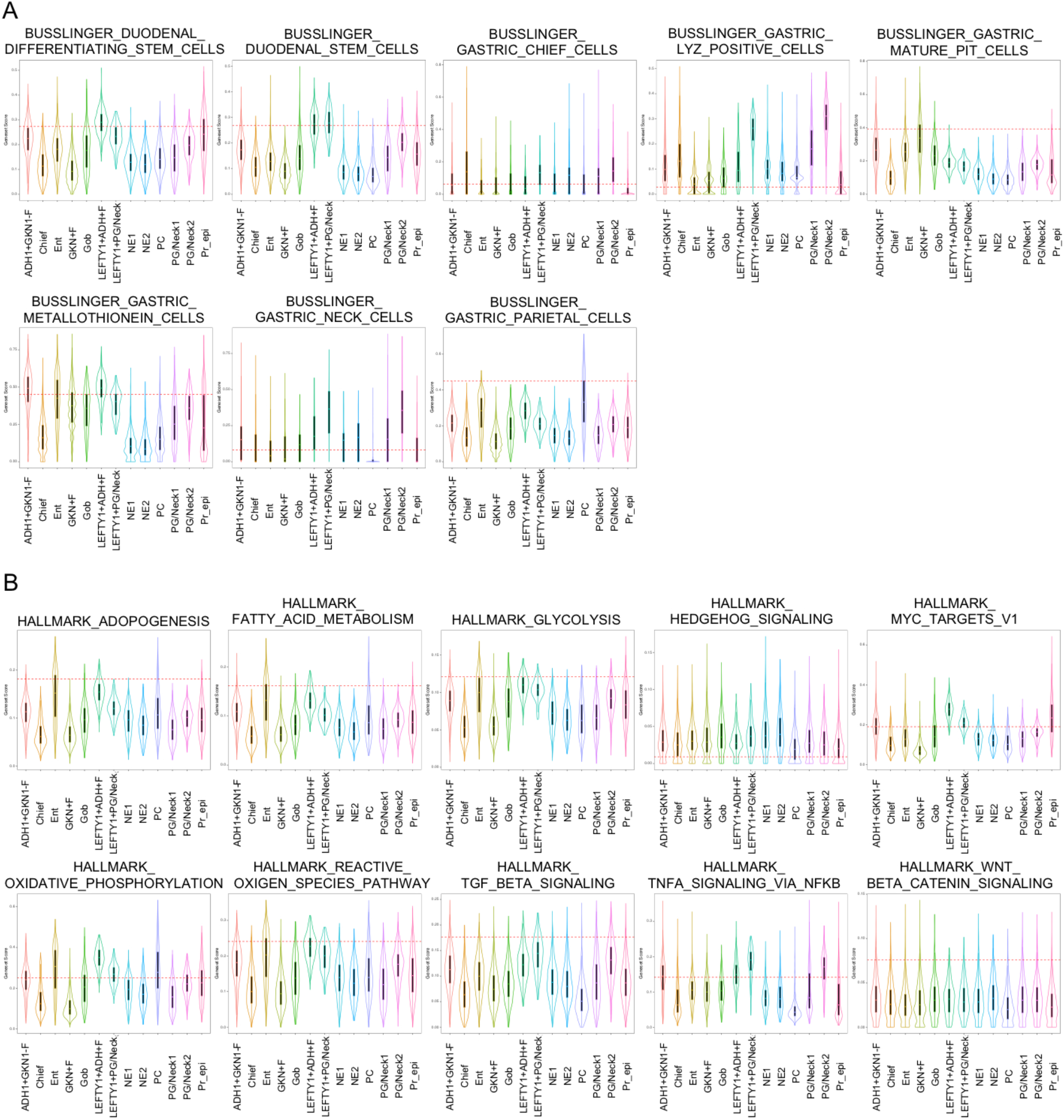
Gene set enrichment analysis of epithelial cells (related to Figures 2–4). (A) Combined violin plots and box plots showing upper gastrointestinal marker gene scores defined by Busslinger et al. (2021). These gene set scores were clearly enriched in our parietal, chief, and neck cell populations, respectively. (B) Combined violin plots and box plots showing HALLMARK pathway scores in each epithelial cell. LEFTY1+ cells show high scores of MYC pathway and metabolic-related gene sets such as adipogenesis, fatty acid metabolism, glycolysis, oxidative phosphorylation, and reactive oxygen species pathways.

**Figure S10.**
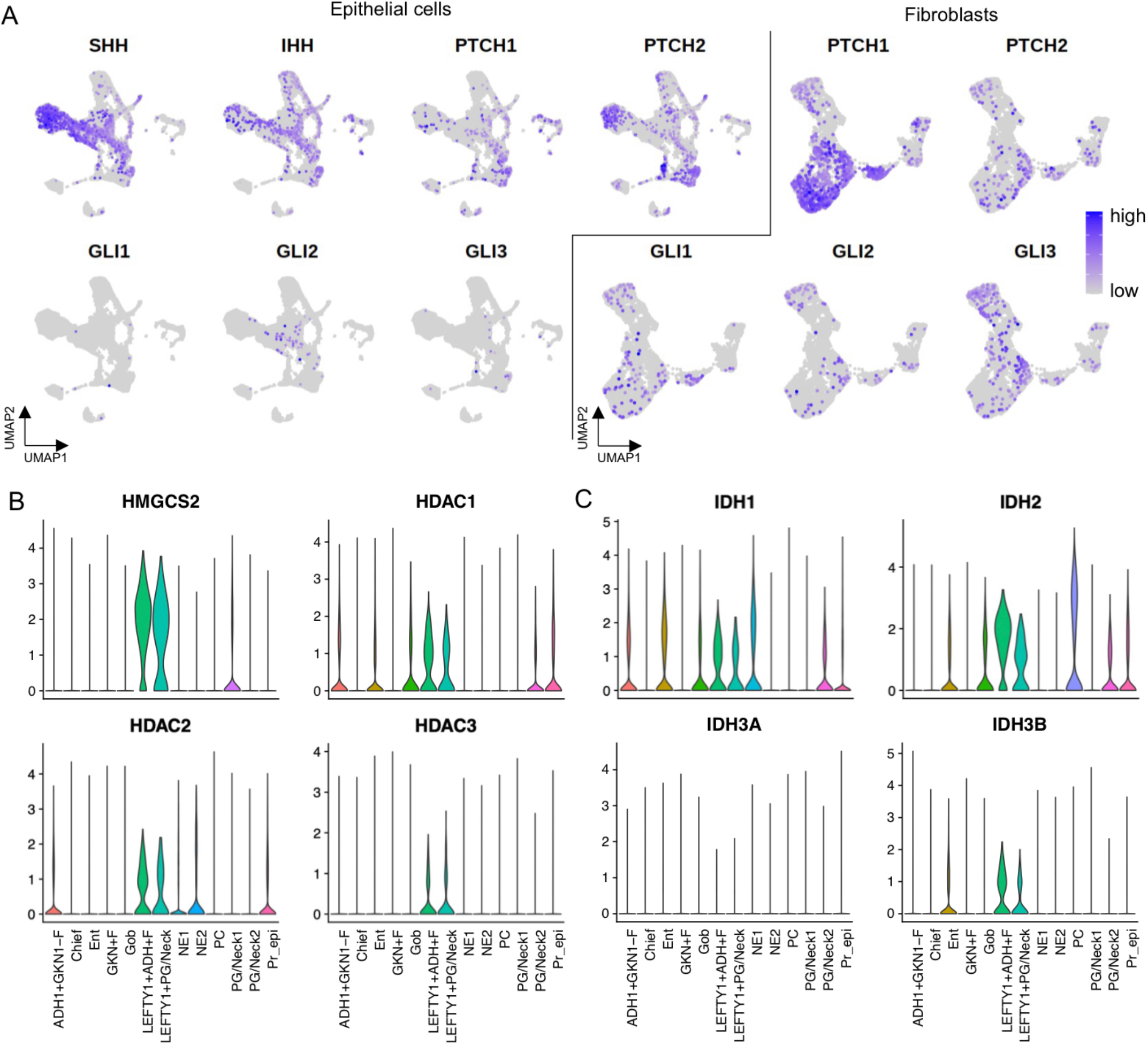
Hedgehog signaling- and metabolite-related gene expression (related to Figure 7). (A) Feature plots showing hedgehog signaling-related genes. SHH expression is limited in gastric lineage, whereas IHH is expressed in both epithelial cells of gastric lineages and metaplastic lineages, respectively. *PTCH1* and *PTCH2* are expressed occasionally in some PG/Neck1 cells and PDGFR+ fibroblasts, and the downstream effectors of Hedgehog, *GLI1*, *GLI2*, and *GLI3*, are expressed modestly in diverse subtypes of epithelial cells and fibroblasts. Left of the black line: epithelial cells; right of the black line: fibroblasts. (B, C) Violin plots showing HMGCS2, HDAC, and IDH expression in epithelial cells. Metabolite-related genes are higher in LEFTY1+ cells.

